# Nanoscale Chromatin Imaging and Analysis (nano-ChIA) platform bridges 4-D chromatin organization with molecular function

**DOI:** 10.1101/2020.01.26.920363

**Authors:** Yue Li, Adam Eshein, Ranya K.A. Virk, Aya Eid, Wenli Wu, Jane Frederick, David VanDerway, Scott Gladstein, Kai Huang, Nicholas M. Anthony, Greta M. Bauer, Xiang Zhou, Vasundhara Agrawal, Emily M. Pujadas, George Esteve, John E. Chandler, Reiner Bleher, Juan J. de Pablo, Igal Szleifer, Vinayak P Dravid, Luay M. Almassalha, Vadim Backman

## Abstract

In eukaryotic cells, chromatin structure is linked to transcription processes through the regulation of genome organization. Extending across multiple length-scales - from the nucleosome to higher-order three-dimensional structures - chromatin is a dynamic system which evolves throughout the lifetime of a cell. However, no individual technique can fully elucidate the behavior of chromatin organization and its relation to molecular function at all length- and timescales at both a single-cell and a cell population level. Herein, we present a multi-technique nanoscale Chromatin Imaging and Analysis (nano-ChIA) platform that bridges electron tomography and optical superresolution imaging of chromatin conformation and transcriptional processes, with resolution down to the level of individual nucleosomes, with high-throughput, label-free analysis of chromatin packing and its dynamics in live cells. Utilizing nano-ChIA, we observed that chromatin is localized into spatially separable packing domains, with an average diameter of around 200 nm, sub-Mb genomic size, and an internal fractal structure. The chromatin packing behavior of these domains is directly influenced by active gene transcription. Furthermore, we demonstrated that the chromatin packing domain structure is correlated among progenitor cells and all their progeny, indicating that the organization of chromatin into fractal packing domains is heritable across cell division. Further studies employing the nano-ChIA platform have the potential to provide a more coherent picture of chromatin structure and its relation to molecular function.

## Introduction

The dynamic, three-dimensional chromatin organization plays an important role in regulating a vast number of cellular processes, including cell-type-specific gene expression and lineage commitment (*1*–*5*). Large-scale alterations in chromatin structure are associated with cancer, numerous neurological and autoimmune disorders, and other complex diseases (*6*–*8*). However, the precise conformation of chromatin and its relationship with transcription, a direct determinant of cellular phenotype, remains contested. The basic units of chromatin are nucleosomes, which are connected by linker DNAs to form a ‘beads-on-a-string’ chromatin fiber. This primary 11-nm fiber was thought to aggregate into thicker a 30-nm chromatin fiber, but this textbook view has been challenged by many studies (*9*–*11*). One of the most recent works includes utilizing a novel imaging technique, chromatin electron tomography (ChromEMT), to interrogate chromatin ultrastructure down to the level of single nucleosomes (*12*). Using ChromEMT, Ou et al. discovered that DNA and nucleosomes assemble into disordered chains, with diameters varying between 5 nm and 24 nm, which themselves pack at various concentrations within the nucleus (*12*).

Parallel to microscopy-based techniques like ChromEMT, chromosome conformation capture techniques (3C, 4C, 5C, and Hi-C) have provided significant insights into higher-order chromatin structures by linking chromatin topology with genomic information (*13*–*16*). Bulk Hi-C measurements, which capture average chromatin structure over millions of cells, have revealed the existence of topologically associating domains (TADs), regions of tens to hundreds of kilobases with frequent intradomain interactions that exhibit a hierarchical organization (*17*–*19*). Notably, single-cell Hi-C methods have demonstrated the potential existence of TADs in individual nuclei, although a high degree of intercellular heterogeneity in TAD distribution has been reported (*20*–*22*).

Recently, the development of super-resolution (SR) microscopy, including stochastic optical reconstruction microscopy (STORM) and photoactivated localization microscopy, in combination with labeling methods, such as fluorescence *in situ* hybridization (FISH) and DNA paint, has brought convergence between microscopy and Hi-C. Multiple independent studies have reported TAD-like chromatin nanocompartments using SR microscopies (*23*, *24*). Additionally, Nozaki et al. elucidated the coherent dynamics of chromatin domains in live cells using SR (PALM) imaging and single-nucleosome tracking (*24*). Despite the advancement in visualizing nanocompartments spatially, there remain several critical open questions, including their packing configuration from chromatin fibers, formation and maintenance mechanisms in live cells, the connection to loci connectivity, and their relationship with transcription processes.

To characterize the details of chromatin organization and understand its relation to the gene transcription at all length scales, it is necessary to overcome several fundamental limitations of current techniques. For Hi-C, the read depth and restriction enzymes limit the resolution to ~1 kb at best, which is insufficient in characterizing individual chromatin chains (*25*). Even SR methods are unable to characterize chromatin structure down to the nucleosome level, as achievable by ChromEMT. However, ChromEMT lacks molecular and genomics information provided by SR and HiC. In addition, ChromEMT, chromatin conformation capture-based, and FISH-derived methods require chemical fixation; thus, only a snapshot of chromatin organization at one point in time can be measured. Consequently, these methods are incapable of monitoring the dynamic process of chromatin reorganization in response to external stimulation and the heritability of higher-order chromatin structure across cell division. Partial Wave Spectroscopic (PWS) microscopy is a label-free, high-throughput spectroscopy technique with live-cell imaging capabilities that have previously been employed to identify and monitor nanoscale structural alterations in chromatin packing in real time (*26*–*28*). Nevertheless, as a diffraction-limited imaging technique, PWS can sense but not resolve chromatin packing at the level of the chain structure. As PWS utilizes the mass-density distribution of chromatin for imaging contrast, it does not carry molecular-specific imaging information. Because no individual technique can fully elucidate the chromatin organization and its relation to molecular function at all length and time scales, it is necessary to develop a multimodal platform combining complementary techniques. Such a platform should possess high resolution across the entire nucleus with dynamic, live-cell imaging capabilities and analysis methodologies to link these results to genome connectivity and the localization of critical molecular factors.

To meet these requirements, we have developed the Nanoscale Chromatin Imaging and Analysis (nano-ChIA) platform, which includes chromatin scanning transmission electron microscopy (ChromSTEM), chromatin transmission electron microscopy (ChromTEM), PWS, and stochastic reconstruction microscopy (STORM). Each facet of nano-ChIA interrogates distinct aspects of chromatin architecture: ChromEM for DNA density and the spatial conformation of the chromatin chains, PWS for label-free, dynamic measurements of the statistical properties of the chromatin conformation in live cells, and STORM for *in situ* imaging of molecular functions with nanoscale resolution. Integrating and coregistering all these modalities, nano-ChIA is a fully quantitative nanoscale imaging platform that complements the genomic information provided by chromatin conformation capture and other sequencing techniques. By bridging high-resolution imaging of chromatin structure and molecular processes with high-throughput, label-free analysis of chromatin dynamics in live cells across timescales, spanning from minutes to hours, nano-ChIA has the potential to provide insights into crucial questions in 3D genomics.

## Results

### nano-ChIA Platform Integrates Information from Multiple Imaging Modalities to Provide Enhanced Spatiotemporal Information on Chromatin Organization and Transcription

The nano-ChIA platform aims to quantify the chromatin organization at broad spatial and temporal scales and relate this structure to transcription activities. At the smallest length scales, the nano-ChIA platform combines DNA-specific labeling (ChromEM) with high-angle annular dark-field (HAADF) imaging in STEM (ChromSTEM) and TEM imaging (ChromTEM). Specifically, ChromSTEM, an adaptation of the pioneering work demonstrated by Ou et al., is able to reconstruct the chromatin structure of a thick nuclear cross-section at sub-3 nm resolution. (Fig. 1A) with the potential to image the entire nucleus by serial sectioning (*29*, *30*). As ChromSTEM is not high throughput (image volume per experiment < 2 μm x 2 μm x 300 nm), the platform utilizes ChromTEM to gain statistical power by imaging ultrathin (< 60 nm) crosssections with larger surface areas (~ 150 μm x 150 um) of different cells within populations. Though not a 3D technique, ChromTEM provides pseudo-2D quantification of the chromatin packing density across the whole nucleus at 3 nm lateral resolution (Fig. 1B). Next, nano-ChIA employs PWS microscopy for label-free, real-time imaging of chromatin packing throughout the nucleus across thousands of cells. PWS directly measures variations in spectral light interference resulting from light scattering due to heterogeneities in chromatin density. This interference signal is then processed to characterize the shape of the autocorrelation function of chromatin density within the coherence volume in both fixed and live cells with sensitivity to structural length scales between 20 and 300 nm (*31*, *32*). To investigate the molecular functionality relevant to chromatin structure, nano-ChIA correlates STORM imaging with PWS on the same cells to visualize chromatin packing structure with respect to the spatial distribution of pertinent macromolecules such as active RNA polymerases (Fig. 1C). A schematic of the combined STORM-PWS microscope is shown in Fig. S1. Finally, nano-ChIA is able to track the timevarying chromatin packing dynamics of single cells using PWS, thus enabling the quantification of supranucleosomal chromatin packing alterations through cell divisions with a temporal resolution on the order of 5 seconds (Fig. 1D).

**Fig. 1.**
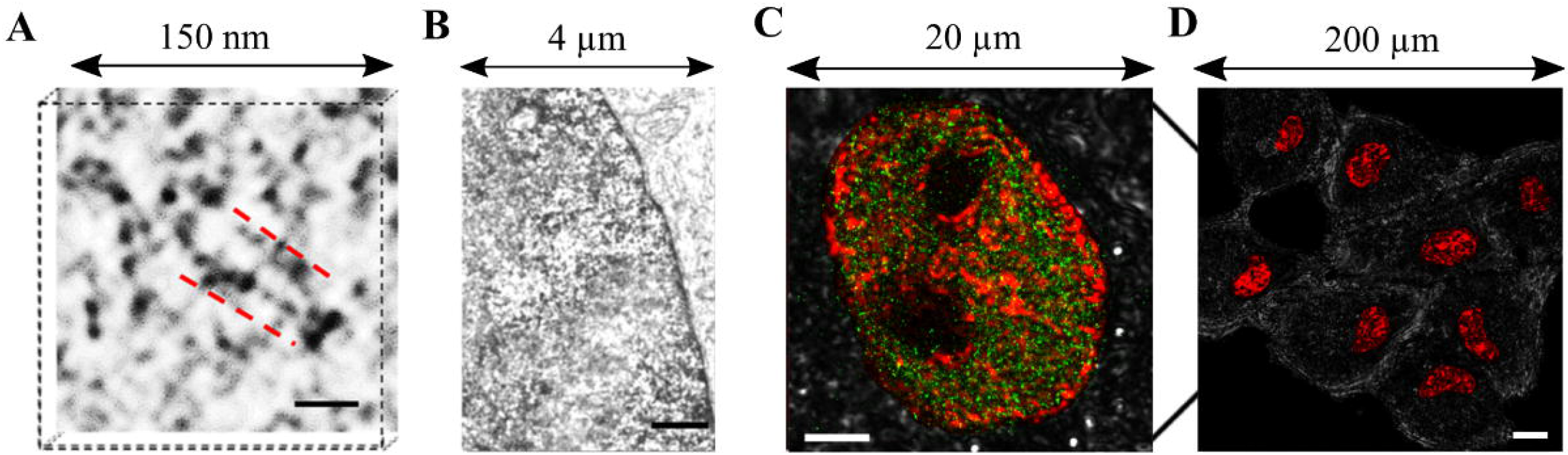
Nanoscale Chromatin Imaging and Nanoscale Analysis (nano-ChIA) platform. (**A**) ChromSTEM HAADF tomography characterizes the 3D chromatin structure of an A549 cell (contrast inverted). The inverted image contrast in ChromSTEM HAADF tomography is inversely proportional to the local DNA density: as the electrons encounter a higher density of DNA along their trajectory, the image contrast appears darker. Individual nucleosomes and linker DNA are resolved at 2 nm spatial resolution. Scale bar: 30 nm. (**B**) ChromTEM imaging of a BJ cell nucleus on a 50 nm resin section prepared by ChromEM staining. Similar to ChromSTEM, ChromTEM also maps the DNA distribution, but the image contrast follows Beer’s law. Scale bar: 1 μm. (**C**) Coregistered PWS and STORM imaging of chromatin packing scaling (*D*, red) and active RNA polymerase II (green) of an M248 cell nucleus. Scale bar: 4 μm. (**D**) Label-free PWS images of live A549 cells. The pseudocolor represents the chromatin packing scaling inside the cell nuclei. Scale bar: 20 μm.

### Chromatin Forms Packing Domains with Fractal Internal Structure

The chromatin polymer adopts a conformation that emerges from interactions between its basic units and the surrounding nucleoplasmic environment, coupled with various biophysical mechanisms that impose additional topological constraints (*33*). A classical homopolymer chain is expected to exhibit self-similar, fractal behavior across all length scales (*34*, *35*). For such a polymer, there exists a power-law mass scaling relation between the number of monomers (*N*) and the size (*r*) of the physical space it occupies: *N* ∝ *r^D^*, where *D* is the fractal dimension or the packing scaling of the polymer (*33*). Assuming that each monomer has an identical molecular weight, the mass of the polymer also scales with the polymer size, following another power-law relationship: *M* ∝ *r^D^*. In a three-dimensional system, fractal behavior occurs for 5/3 < *D* < 3 depending on the balance of the free energy of polymer-polymer interactions versus the free energy of polymer-solvent interactions. For an ideal chain in a theta solvent, where the free energy of monomer-monomer interactions and the free energy of monomer-solvent interactions are equally preferred, the fractal dimension *D* = 2. In a good solvent, the free energy of monomer-solvent interactions exceeds that of monomer-monomer interactions and 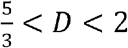. In contrast, monomer-monomer interactions are preferred in poor solvent conditions, leading to polymer collapse and 2 < *D* < 3 if the fractal structure is preserved, which is not guaranteed. However, unequivocal evidence regarding whether chromatin exhibits fractal behavior and at what length scales this behavior occurs is still absent. To accomplish this task, we map the relationship between the genomic length of chromatin and the physical space it occupies by leveraging the sub-3 nm spatial resolution of ChromSTEM for fixed cells and the nanoscopic sensitivity of PWS for live cells.

First, we utilized ChromSTEM in nano-ChIA to reconstruct high-resolution 3D chromatin structure with a resolution below 3 nm from part of the nuclei of four A549 lung adenocarcinoma cells (Fig. 2A). ChromSTEM was able to resolve variably packed individual nucleosome assemblies with clear linker DNA segments (Fig. 2B-2C). We utilized mass scaling analysis to quantify how chromatin mass (*M*) contained within volume *V* scales up with the radius *r* of that volume 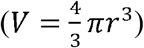. Experimentally, we randomly sampled different regions of the chromatin and calculated an average mass scaling curve. Two power-law regimes were observed on the mass scaling curve, each with a distinct scaling exponent. The power scaling exponent of the two regimes were calculated by two linear regression lines to the mass scaling curve in log-log scale: fit to the first and last 50 nm range on the mass scaling curve respectively. In principle, there should exist a third region from 0 nm to 12 nm, which represent the chromatin packing within the chromatin chains (2.5 nm to 12 nm in radius) as first observed in earlier ChromEMT work (*12*), we suspect that the resolution of ChromSTEM (~ 2 nm) is not sensitive enough to fully probe such fine strucutre. The scaling exponent of each of these power-law regimes was estimated by performing linear regression on a log-log scale. The first regime has a scaling exponent, *D*=2.58, which indicates that the chromatin packing in this regime adopts a fractal structure (*D* between 5/3 and 3). For the second regime, the scaling exponent increases to 3.02, indicating a switch to a structure that is almost completely space-filling. There is a smooth transition between the fractal regime and the space-filling regime, as the slope of the mass scaling curve on a log-log scale gradually increases at the boundary of the fractal regime. We hypothesized that individual domains with internal fractal structures exist. Each of these domains likely has different packing scaling behavior, *D* and domain size. Consequently, the average mass scaling curve, calculated by the superposition of the mass scaling curves for each domain, can show such a transition.

**Fig. 2.**
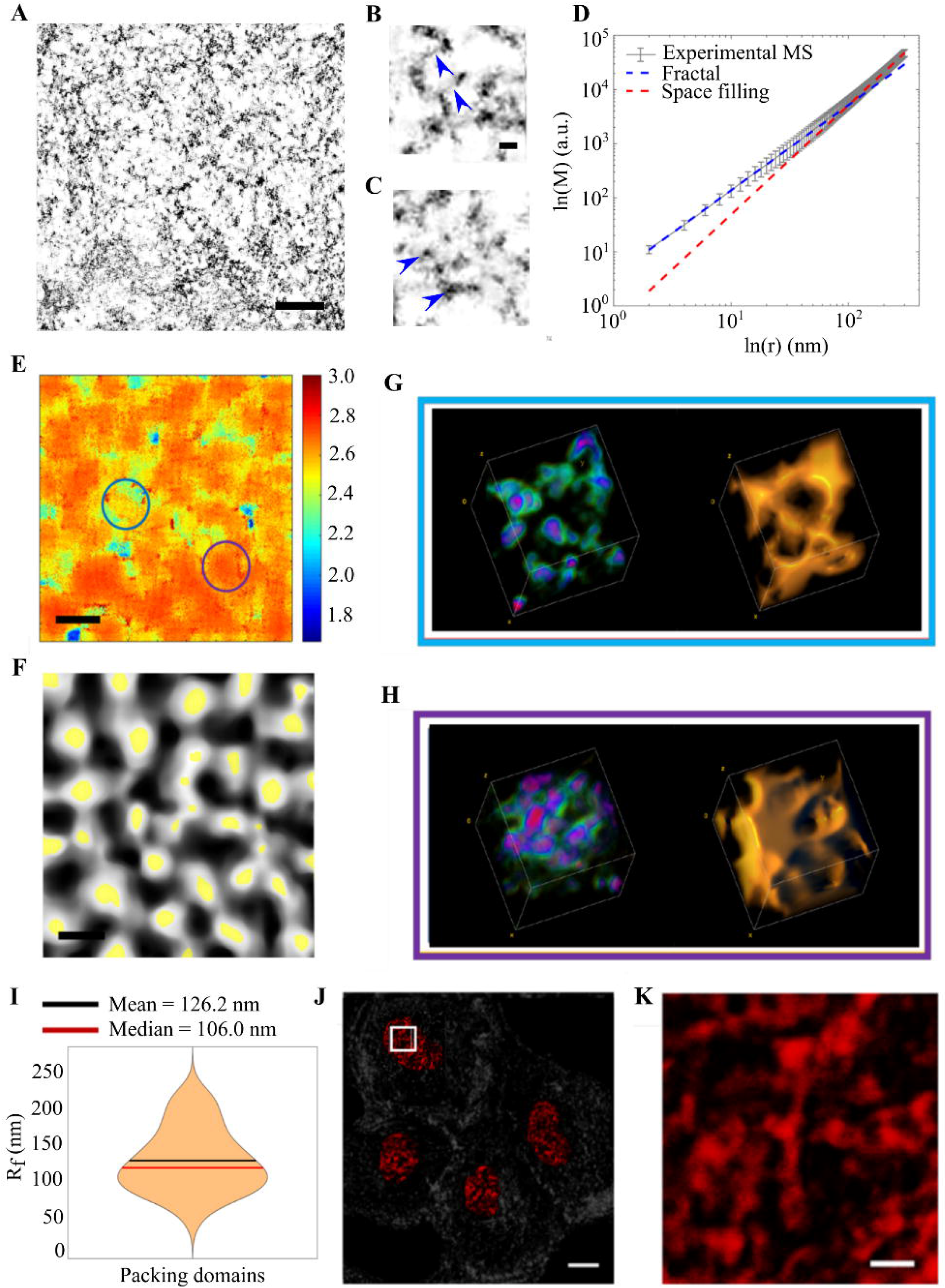
nano-ChIA identifies chromatin packing domains (PDs). (**A**) A 2.9 nm thick virtual 2D slice from ChromSTEM HAADF tomography reconstruction of chromatin from an A549 cell nucleus (contrast inverted). Scale bar: 200 nm. (**B-C**) High-resolution tomography reveals fine chromatin structures such as linker DNA, as shown by blue arrows in (**B**), and individual nucleosomes, as shown by blue arrows in (**C**). Scale bar: 20 nm. (**D**) Average mass scaling of the chromatin packing in the log-log scale shows a packing domain regime with an internal fractal structure. Within individual PDs, genomic distance and the physical space occupied by chromatin follow a power-law relationship (linear in log-log plot). The dashed lines depict the linear regression fit within each regime. (**E**) Mapping of chromatin packing scaling (*D*) of an A549 cell. The chromatin self-organizes into packing domains (PDs) with similar packing scaling. Scale bar: 300 nm. (**F**) Segmentation of *D* mapping. White regions represent the PDs, and yellow regions depict the center of these PDs determined by the flooding algorithm. Scale bar: 300 nm. (**G-H**) Supranucleosomal packing configurations for two PDs with different *D*s in (**F**): (g) corresponds to the area highlighted by the blue arrow in (**F**), and (**H**) corresponds to the purple arrow in (**F**). In the leftmost rendering of each panel, the DNA concentration increases from green to red. The rightmost rendering shows the surface topology of the two PDs. (**I**) Distribution of the radii of the PDs (R_*f*_). The mean and median radius of the PDs corresponds to the upper bound of the fractal regime of the mass scaling curve. (**J**) *D* mapping of the whole cell by PWS. Scale bar: 10 μm. (k) *D* mapping by PWS. Each cluster represents a diffraction-limited observation of PDs. Scale bar: 1 μm.

Next, we mapped the distribution of the chromatin packing scaling *D* measured from the fractal regions of the mass scaling curve for the entire field of view (Fig. 2E). The *D* distribution showed that chromatin organizes into spatially separable fractal packing domains (PDs) with similar packing scaling. From the *D* distribution, we identified the center of each domain by employing a flooding segmentation algorithm (Fig. 2F). We estimated the radius of fractal packing domains (*R_f_*) as the distance at which the single mass scaling curve for the domain significantly deviates from a power-law relationship, the details of the algorithm can be found in SI. We observed that *R_f_* has a median value of 106.0 nm, which agrees qualitatively with the upper bound of the fractal regime calculated from the average mass scaling curve (Fig. 2I). The existence of PDs with various *D* and *R_f_* can provide an explanation for the gradual transition between the fractal and space-filling regimes. Notably, a low-*D* domain (Fig. 2E blue circle, Fig. 2G) exhibits a profoundly different supranucleosomal packing configuration from a high-*D* domain (Fig 2E purple circle, Fig. 2H).

As ChromSTEM has a limited field of view and requires chemical fixation, we employed PWS to inspect the chromatin packing scaling distribution across the entire nucleus and confirm the presence of packing domains in live cells. As previously mentioned, PWS measures chromatin density fluctuations. Chromatin packing scaling, *D*, can be calculated from these fluctuations, as explained (Section S1). In agreement with ChromSTEM, PWS analysis also identified spatially separable chromatin packing domains characterized by similar D values (Fig. 2J-2K). In summary, by combining ChromSTEM and PWS, we have identified the existence of supranucleosomal chromatin packing domains within which the chromatin chain exhibits fractal behavior. A natural next step is to investigate the potential functional significance of these domains.

### Relationship Between Chromatin Packing and Genome Connectivity

Besides packing scaling, contact probability scaling is another important statistical property of chromatin that represents overall polymer connectivity and can be measured using chromatin capture techniques such as Hi-C. For our classical polymeric systems, the probability of contact (*P*) between two monomers of linear distance *r* apart also follows a power-law scaling relation: *P* ∝ *N^−s^*, where *s* is the contact probability scaling. For a random coil in a theta solvent, *s* = 3/2, while for a fractal globule in a poor solvent, *s* = 1(*36*). Recent advances in Hi-C and super resolution imaging techniques have demonstrated that no single power-law scaling exponent can describe chromatin organization throughout the nucleus, and several studies have disproved the contact probability scaling behavior of chromatin predicted by the fractal globule model (*17*–*19*). Classical homopolymers exhibit an inverse relationship between contact probability scaling and fractal dimension. For example, both the ideal chain and fractal globule models have a simple analytical relationship between these two properties, specifically *s* = 3/*D*(*36*). However, the relationship between chromatin packing behavior and genome connectivity has not been investigated in fractal chromatin PDs.

Because no existing model can faithfully capture all aspects of chromatin structure, we sought to test this hypothesis by implementing two distinct models of chromatin. The models we employ here are not expected to be an exhaustive set but instead were used as testbeds to ascertain whether the inverse relationship between *s* and *D* was likely to be a model-independent property. First, we implemented a basic homopolymer model to represent chromatin structure within packing domains. We introduced effective attractions between monomers using the classic Lennard-Jones potential, which physiologically represents the solvent quality of the polymer solution. We tuned the attractive potential between monomers to generate polymers ranging from a swollen self-avoiding walk coil to a collapsed globule in order to modulate two measurable statistical polymeric properties, *D* and *s*, and investigated their relationship (*17*). Additionally, we employed the self-returning random walk (SRRW) model that implements a heterogeneous distribution of monomers to represent the high degree of conformational freedom of the disordered chromatin polymer (*37*). Importantly, the model explicitly includes stochastic, self-returning events to create TAD-like structures with frequent self-contacts. Each tree-like topological domain is connected and isolated by open backbone segments. The probability of self-returning events is controlled by the chromatin folding parameter, which modulates the size and packing behavior of hierarchical tree domains. We tuned this parameter to modulate the fractal dimension and contact probability scaling to evaluate their relationship for this particular chromatin model (*37*). SRRW conformations were generated by Monte Carlo (MC) simulations with each step size representing 2 kb of DNA (~10 nucleosomes). For both models, we measured *D* and *s* by performing a linear regression on their respective power-law scaling relations. The regression was performed within the genomic range from 20 kb to 200 kb, which is of the same order of magnitude as the size of chromatin PDs. Although the two chromatin models resulted in two distinct functional forms of *s*(*D*), as would be expected, both models demonstrated an inverse relationship between these two statistical parameters (Fig. 3A-3B). This result suggests that the inverse relationship between *D* and *s* could be a general model-independent property of self-similar polymer domains. After computationally establishing the inverse relationship between space-filling behavior and polymer connectivity, we wanted to investigate whether this property can be observed *in vitro*.

**Fig. 3.**
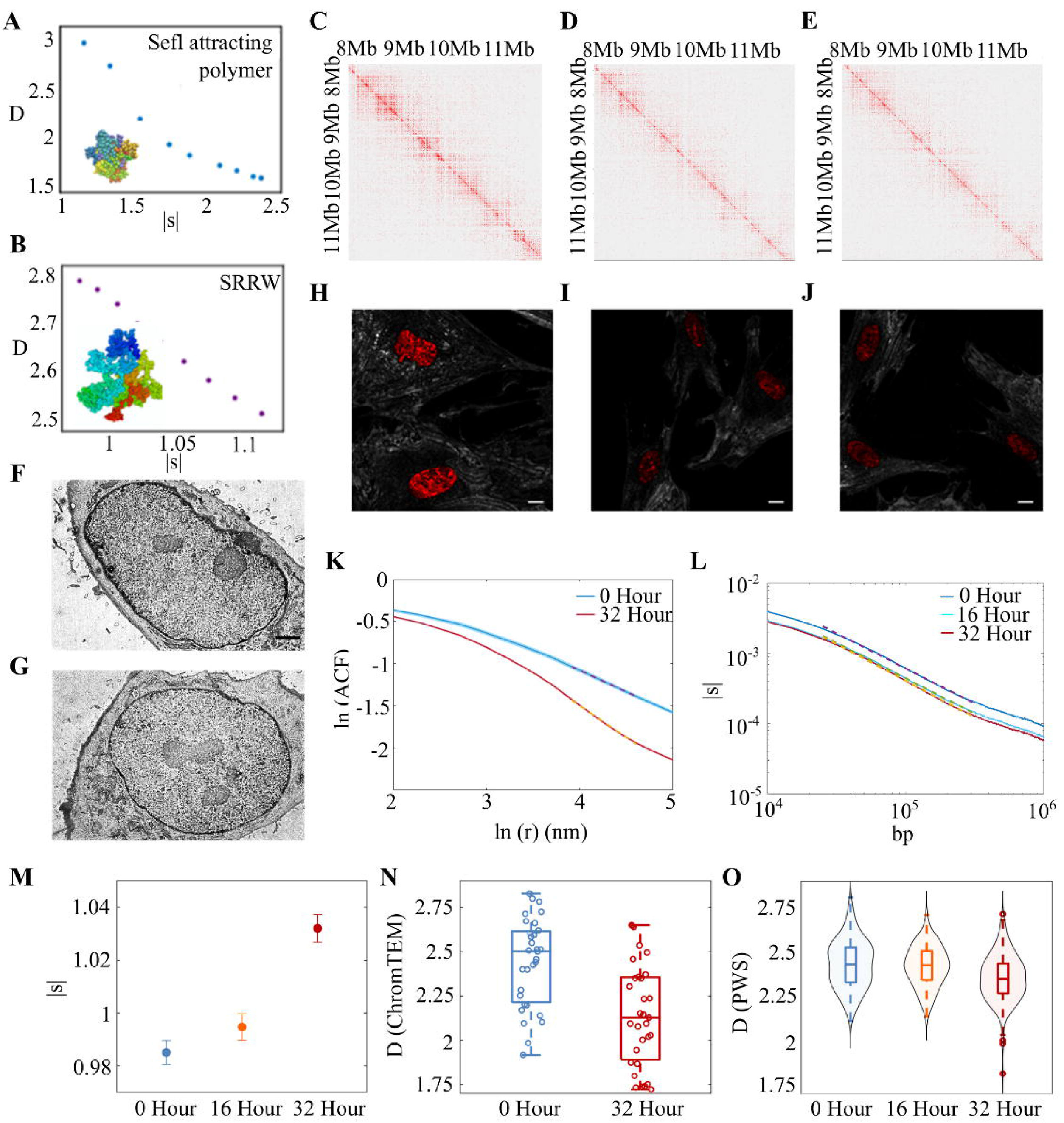
Measuring chromatin packing scaling alterations induced by dexamethasone (DXM) in BJ cells. Computational models reveal a general inverse relationship between *s* and *D* using selfattracting polymer (**A**) and self-returning random walk (**B**). We visualized the contact probability maps from publicly available whole genome intrachromosomal Hi-C data for BJ cells with DXM treatment at 0-hour (**C**), 16-hour (**D**), and 32-hour (**E**) time points. ChromTEM images of BJ cells without DXM treatment (**F**) and with 32-hour (**G**) treatment. Scale bar: 2 μm. (**H**-**J**) PWS images of BJ cell with DXM treatment at 0-hour, 16-hour, and 32-hour time points, respectively. Scale bar: 10 μm. (**K**) ACF analysis using ChromTEM. The average ACF of the control group (blue) is significantly different from the average ACF of the treated group (red). *D* was measured inside the first fractal domain (50 nm to 100nm) by a linear regression fit of the ACF in the log-log scale. (**L**) Contact probability scaling *s* for BJ cells treated with dexamethasone for 0 hours, 16 hours, and 32 hours, calculated from intrachromosomal Hi-C contact data. The linear regression fit was performed on contact probability versus genomic distance between 10^5.8^ and 10^6.8^ bp. (**M**–**O**) Chromatin packing scaling alterations induced by DXM treatment measured from Hi-C contact data, ACF analysis of ChromTEM images, and PWS. Consistent changes were observed across the nano-ChIA platform.

To test this hypothesis experimentally, we employed the nano-ChIA platform to measure changes in chromatin packing scaling *D* upon external stimulation, which we compared with changes in contact probability scaling *s* measured by Hi-C analysis. Dexamethasone (DXM) treatment has previously been demonstrated to alter whole-scale genome connectivity (*38*). Analysis of publicly available Hi-C data revealed that *s* increases at 16 hour and 32 hour time points in BJ cells (Human fibroblast cells) treated with 100 nm DXM (Fig. 3C-3E, 3L-3M), which we hypothesized would result in an inverse change in chromatin packing scaling. Thus, using ChromTEM (Fig. 3F-3G) and PWS microscopy (Fig. 3H-3J), we measured changes in *D* before and after treatment with DXM in fixed and live cells. Unlike ChromSTEM, which resolves exact 3D structure, ChromTEM images the projection of a thin cross section (50 nm) of chromatin. To calculate chromatin packing scaling *D* from ChromTEM data, we utilized autocorrelation function (ACF) analysis (Fig. 3K). The ACF of the spatial variations of the density of a polymer, such as chromatin, can be derived from its mass scaling relationship: 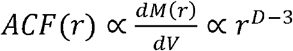, where *M*(*r*) is the mass of chromatin contained within volume 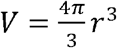. Through this relationship, *D* can be calculated from the power-law scaling exponent of the ACF. For an infinite, continuous, random structure, the 2D ACF of the projection of a thin section can be considered to be identical to the 3D ACF of the original 3D structure with high accuracy. For a finite fractal structure, we showed numerically that the *D* estimated from the 2D ACF of a 50 nm thin section introduces less than 3% error (Fig. S2).

In agreement with our modeling results, we indeed observed inverse changes in *D* and *s* at the level of individual cells, as measured by both ChromTEM and live-cell PWS microscopy (Fig. 3N-3O). Notably, we demonstrated the ability of ChromTEM and PWS microscopy to acquire information in agreement with each other and complementary to bulk chromosome conformation capture methods. Importantly, live-cell PWS microscopy can measure real-time changes in chromatin packing scaling without the need for fixation, digestion, or fluorescent labeling and with the potential for high-throughput imaging. To demonstrate the live-cell imaging capabilities of PWS, we tracked a subset of BJ cells after DXM treatment, quantifying chromatin structure every 4 hours for a total of 32 hours (Fig. S5). Additionally, we employed PWS to track the same cell over 32 hours to visualize alterations in chromatin structure dynamically (Fig. S6). Neither Hi-C nor ChromTEM possesses these real-time monitoring abilities, as both techniques require chemical fixation and thus provide only a snapshot of chromatin structure at one point in time.

To further test the inverse relationship between *D* and *s*, we performed additional PWS experiments with A549 cells treated with DXM for 12 hours, and compared the results to publicly available Hi-C results(*39*). Again, we observed the same inverse relationship between *D* and *s* (Fig. S3, Fig. S4). Altogether, these results suggest that, *in vitro*, genome connectivity is related to the fractal behavior of chromatin within PDs, in agreement with the behavior predicted from our simulations. Additionally, we have demonstrated that nano-ChIA allows real-time monitoring of chromatin structure and associated statistical properties of genome connectivity in live cells. Finally, nano-ChIA can be used in the future to establish the exact relationship between chromatin packing and genome connectivity inside packing domains, which in turn can help test and identify an optimal model of chromatin structure.

### Relationship Between Chromatin Packing and Transcription

Next, we wanted to leverage nano-ChIA to explore the relationship between chromatin packing and gene transcription. Earlier studies have suggested that chromatin density may play a role in the regulation of transcriptional processes (*24*, *27*, *40*, *41*). To predict large-scale patterns of gene transcription through the chromatin packing scaling *D*, we have developed a multiscale computational chromatin packing macromolecular crowding (CPMC) model that combines the crowding effects with the effects of other molecular and physical regulators of transcription. As *D* of a fractal PD increases, the model predicts increases in the accessible surface area of chromatin as well as the variance in crowding conditions to which the genes within the domain are exposed. In turn, chromatin density within a given transcriptional interaction volume increases the rates of binding of transcriptional reactants and decreases their rates of diffusion. As a result, at lower *D*, gene expression will asymptotically increase with *D* up to an inflection point. At this critical value of specific chromatin packing scaling, the range of crowding conditions to which the genes are exposed becomes suboptimal. Thus, after a certain critical *D* value, the transcriptional output is predicted to decrease. The shape of this nonmonotonic relationship between *D* and gene expression is dependent on several molecular and physical regulators of transcription (defined in **Table S1**). For example, higher concentrations of transcriptional reactants increase ensemble expression rates across all *D* values (Fig. 4A). Additionally, these more favorable molecular conditions shift the critical chromatin packing scaling to higher values.

**Fig. 4.**
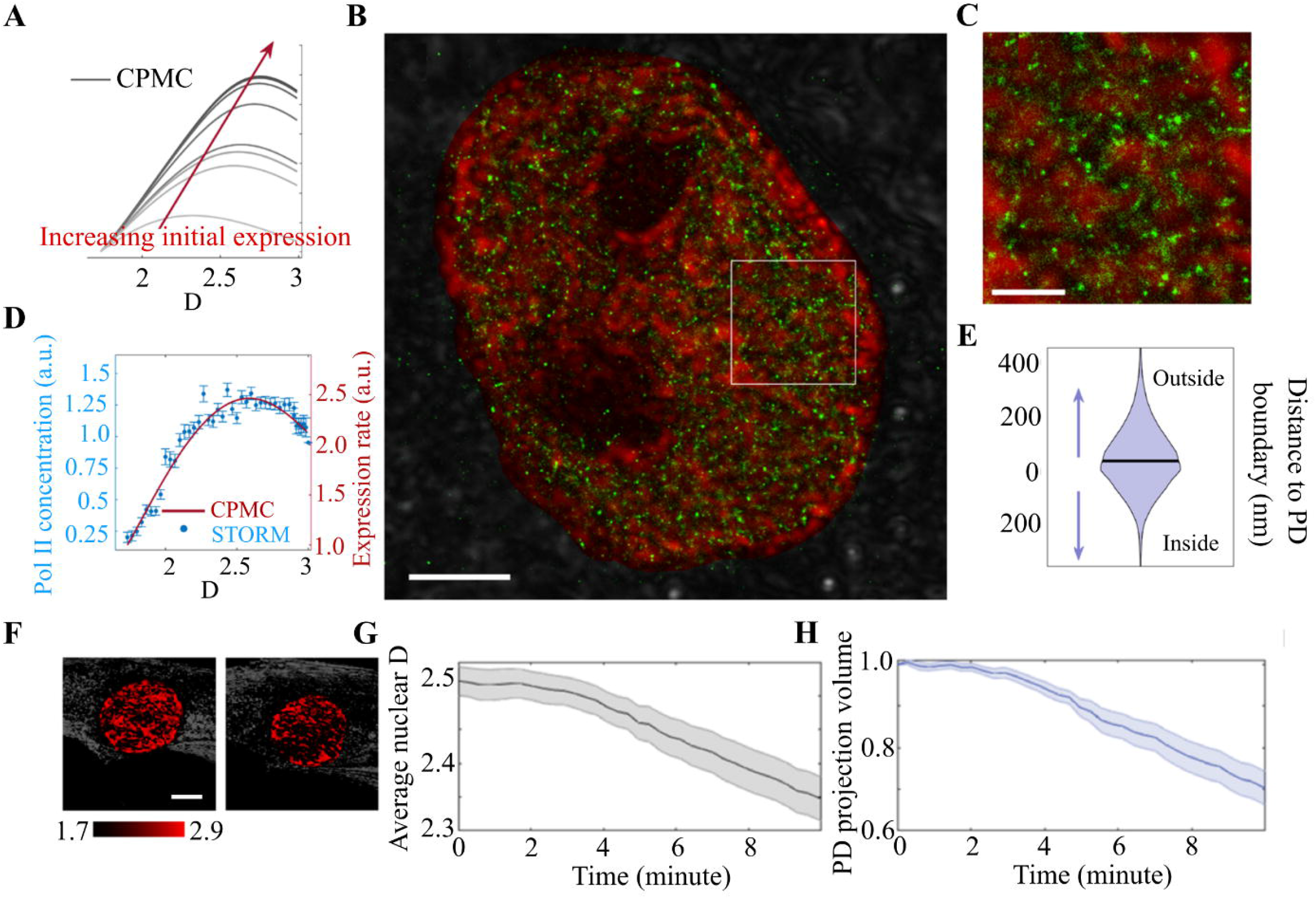
Nano-ChIA platform investigates the relationship between chromatin structure and transcription. (**A**) Multiple implementations of the CPMC model for low- and high-expression genes show that in all cases, the surrounding chromatin packing scaling has a nonmonotonic relationship with gene expression. (**B**) A STORM image of an M248 ovarian cancer cell labeled with Pol II (green) overlaid on top of chromatin packing scaling D measured by PWS (red). Scale bar: 3 μm. (**C**) A magnified view of the white square in (**A**). Scale bar: 1 μm. (**D**) The relationship between *D* (chromatin packing scaling) and the local concentration of Pol II (gene expression level) compared with one realization of the CPMC model. (**E**) A violin plot shows the distribution of distances between enriched Pol II regions and their nearest packing domain group. The plot shows that Pol II tends to distribute around the boundary of packing domains. (**F**) PWS imaging of a live BJ fibroblast cell during Act-D treatment. Scale bar: 5 μm. (**G**) After transcriptional elongation is halted with Act-D, average nuclear chromatin packing scaling decreases steadily in a matter of minutes as measured by PWS. (**H**) The change in the volume fraction of the nucleus containing packing domains as measured by PWS.

Experimentally, we employed STORM-PWS colocalization to investigate the relationship between chromatin structure and gene expression. We utilized the STORM-PWS to localize regions of active gene transcription by imaging active RNA Polymerase-II (Pol-II) with STORM and measured the surrounding chromatin packing scaling through PWS (Fig. 4B-4C). Trends predicted by the CPMC model were in excellent agreement with the *in situ* experimental STORM-PWS findings across multiple cell lines, demonstrating a consistent nonmonotonic relationship between chromatin packing structure (*D*) and transcription (Fig. 4D, Fig. S8). Notably, we found that Pol-II density is associated with observed PDs, with pockets of high Pol-II density forming around the periphery of these high-*D* structures (Fig. 4A-4C). Additionally, these pockets of higher Pol-II density appear to occur primarily on the periphery of larger groups of PDs or on the periphery of smaller, isolated PDs.

The agreement between the CPMC model and the experimental data seems to support that the chromatin packing structure can influence the rate of transcription. However, these results do not exclude the possibility that transcription reactions will affect chromatin packing, suggested by the preferred localization of Pol-II at the periphery of PDs. In order to test the second hypothesis, we performed a perturbation study to halt transcriptional elongation in BJ fibroblast cells by treatment with Actinomycin D (Act-D). After Act-D treatment, we continuously captured PWS images for ten minutes to evaluate the real-time effect of transcriptional inhibition on chromatin structure from the level of PDs to the scale of the whole nucleus (*42*, *43*). For each PD, PWS measured *D* with sensitivity down to the size of the chromatin chain (20 nm)(*32*). At the level of the entire nucleus, we observed that treatment with Act-D produces a rapid decrease in average chromatin packing scaling across the cell population (Fig 4G). Over ten minutes, we observed a 7% decrease in average nuclear *D*. Additionally, we observed that the projection fraction of the nucleus consisting of packing domains, measured by PWS, decreased by 29% (Fig 4H). Importantly, the abrogation of transcription did not eliminate the PD structure of chromatin (Fig. S11).

Our findings are consistent with previous work showing that chromatin structure is stabilized by transcriptional elongation (*19*, *24*, *44*). Furthermore, this result supports the hypothesis that the process of active gene transcription affects supranucleosomal chromatin organization but is not its sole determinant. In conclusion, our interrogation of the relationship between chromatin structure and transcription provides evidence of a complicated, bidirectional relationship between supranucleosomal chromatin packing and gene expression.

### Chromatin Packing Domains are Heritable Across Cell Division

Cell-type-specific gene expression patterns are inherited through mitosis. A hierarchy of gene expression patterns is re-established after mitotic exit to ensure the maintenance of cell identity, potentially driven by mechanisms such as mitotic bookmarking through the maintenance of histone modifications at promoter regions (*45*–*47*). Additional studies using chromosome conformation capture techniques have demonstrated that higher-order cell-type-specific structures, such as TADs, are lost during mitosis and re-established along with a lineage-specific replication timing program in early G1 (*48*, *49*). Altogether, these results suggest a potential relationship between transcriptional memory propagation and the heritability of chromatin structure. These considerations led us to employ live-cell PWS from our nano-ChIA platform to investigate whether chromatin packing behavior is transferred between parent and progeny cells through cell division. Critical questions include how the spatial distribution of the scaling of chromatin packing evolves over long timescales (hours) and whether the time-dependent fluctuations of the chromatin packing scaling across the nucleus are conserved through the process of cell division.

To address these questions, we measured chromatin packing scaling in HCT116 colon cancer cells with PWS for 20 hours. During that time, several cell division events were observed (Fig. 5, SI Video 1, 2). We isolated ten dividing cells and calculated a histogram showing the spread of *D* across each cell. We then calculated the ratio of each histogram by the average histogram for all cells at each timepoint (relative to that measured at the time point of cell division), thus focusing on each cell’s unique deviation from the population mean (see Fig. S12 for step by step analysis). After cell division, we compared the histogram ratios of cells that originated from the same progenitor and cells originating from unique progenitors, as shown in Fig. 5A-5B. We found that related progeny cells are more highly correlated with each other over time, even several hours after cell division. We also compared the histogram ratio from each progeny cell 3 hours after cell division to the histogram ratio of all progenitor cells 3 hours before cell division. We also found a significantly higher correlation between progeny cells and their related progenitors than between progeny cells and unrelated progenitors in the same population (Fig. 5C).

**Fig. 5.**
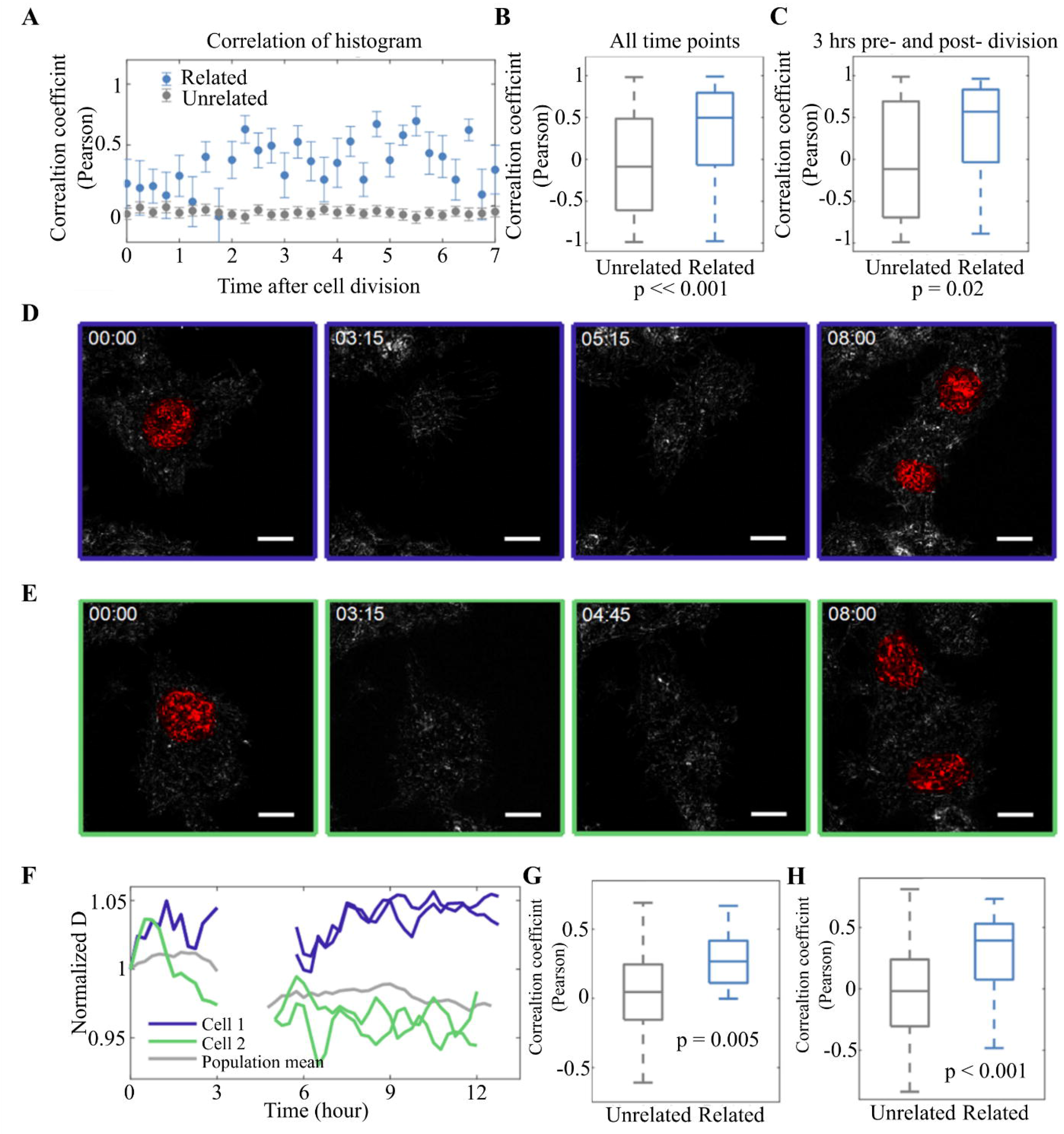
Time-resolved imaging of chromatin packing scaling during cell division in HCT115 COLON CANCE. (**A**) After cell division, the normalized histograms of paired progeny cells are more highly correlated with each other than other progeny cells at the same time point. (**B**) Across all time points, there is a highly significant correlation showing that paired progeny cells are more highly correlated with each other than with other cells. (**C**) Comparing all progeny cells 3 hours after cell division to all progenitors 3 hours before cell division shows that progeny cells have a higher correlation with their “parent” progenitor cell than with unrelated progenitor cells. (**D**-**E**) PWS images showing chromatin packing scaling at four time points of the dividing cells. During cell division, the cells lift slightly off the glass, causing them to exit the depth of field of the objective. When they return to the glass, they have divided. (**F**) Average nuclear D from the cells shown in (**A**) and (**B**) are tracked over time. After ~5 hours, both cells had divided, and their progeny cells were tracked for ~7 hours. (**G**) Progeny cells are more strongly correlated with their paired progeny cells than with other unrelated cells on the dish (N = 20 progeny cells). (**H**) Progeny cells appear to be more correlated with their progenitor cells than with other unrelated cells on the dish (N = 20 progeny cells).

To track how the average chromatin packing scaling might change over time, we tracked the mean packing scaling for all cells (n = 10). Fig. 5D-5E show the tracking of two dividing cells over time. We observe that the average nuclear *D* has relatively small fluctuations compared to the mean before cell division, which is uncorrelated from cell to cell (Fig. 5F). During division, the cells lift up and partially detach from the dish, causing the nucleus to exit the detectable field of view of the PWS system. Following adherence to the glass, the progeny of the original cells reappear with a resumption of small fluctuations in chromatin packing scaling. Notably, the time-dependent changes in the chromatin packing scaling of each pair of progeny cells were more likely to be correlated with each other than with other dividing cells (Fig. 5G). Likewise, the chromatin packing scaling of progeny cells during the two hours following mitosis is more correlated with their progenitors than with other cells at the same temporal cross-section following division (Fig. 5H). Altogether, these results demonstrate that the chromatin packing domain structure evolves and is heritable through the process of cell division.

## Discussion

In this work, we present nano-ChIA, a multi-technique nanoscale imaging and analysis platform that enables the study of chromatin organization across a wide range of length and time scales (Fig. 1). At sub-3 nm spatial resolution, ChromSTEM provides three-dimensional information about chromatin configuration down to the individual nucleosome and DNA strand level. However, the ChromEM staining used in both TEM imaging and ChromSTEM tomography requires chemical fixation and can only measure chromatin organization at a single time point. A complementary nanoimaging technique, PWS, provides a label-free imaging method with livecell capabilities that enables probing the dynamics of chromatin structure in individual cells, in real-time, with a subsecond temporal resolution, and across entire cell populations. Although diffraction-limited, PWS is sensitive to chromatin packing in a range from 20 nm to 300 nm (*32*). ChromSTEM and PWS provide the physical context of chromatin structure, i.e., they quantify chromatin conformation. The third component of nano-ChIA, STORM, coregisters the locations of targeted molecular species or events with 20 nm localization precision, thus providing the molecular functionality of chromatin organization. Collectively, the nano-ChIA system integrates structural information about chromatin packing with details regarding the localization of critical molecular factors.

Consolidating results from electron and PWS microscopy allowed us to uncover the existence of chromatin packing domains (PDs) with internal fractal structures of varying sizes throughout the nucleus in both fixed and live cells (Fig. 2). Additionally, we determined that contact probability scaling and chromatin packing scaling within these packing domains follow an inverse relationship through both polymer simulations and experimental cross-validation with Hi-C (Fig. 3). This suggests that the physical packing of chromatin into fractal domains may affect contacts between genes located within the same domains. Employing STORM molecular nanoscopy, we were able to interrogate the relationship between chromatin structure and transcription processes (Fig. 4). We found that the chromatin packing scaling of a domain influences the extent of active transcription within the PD. Conversely, these transcription processes themselves can also influence the organization of chromatin packing domains, as the disruption of transcriptional elongation results in significant changes in the packing scaling of these domains. Finally, we exploited the dynamics capabilities of PWS to assess the heritability of supranucleosomal chromatin organization between progenitor and progeny cells through the process of cell division. We demonstrate that chromatin packing scaling is correlated among progeny cells from the same progenitor as well as between progeny cells and their progenitors (Fig. 5).

The supranucleosomal chromatin structure uncovered by nano-ChIA suggests that chromatin fibers (‘beads-on-a-string’) can pack into spatially separable PDs of varying genomic sizes and densities. Importantly, we observe a power-law mass scaling behavior of chromatin conformation inside the packing domains, indicating that chromatin adopts locally self-similar fractal structures. The existence of fractal packing domains illustrates the statistical behavior of chromatin as a polymer. Chromatin may adopt a variety of distinct configurations in 3D space but still produce the same statistical chromatin packing behavior, which can be quantified by *D*. Specifically, the exact chromatin structure surrounding a specific genomic locus within a given PD may differ across a cell population and over time. However, the statistical properties of the encompassing PDs, such as their size and packing scaling, may remain the same.

Packing structures can be formed by a variety of mechanisms – from changing polymer-polymer and polymer-solvent interactions, as during phase separation, to the introduction of topological constraints, such as confining chromatin loops by CTCF/cohesin complexes, transcription-induced supercoiling, or interactions with lamin (*50*–*58*). In particular, polymers with the higher free energy of self-interactions than of polymer-solvent interactions tend to adopt conformations with a higher fractal dimension. In turn, self-interactions and chromatin-nucleoplasm interactions might be affected by chromatin chain and nucleoplasm nanoenvironment modifications, such as histone modifications, DNA methylation, nucleoplasmic crowding, pH, and ionic environment. The presence of chromatin PDs with distinct fractal behavior contradicts the prediction that all chromatin within the nucleus forms a single, self-similar structure and is thus discordant with the view of chromatin as a homopolymer. Intuitively, the heteropolymeric behavior of chromatin is consistent with what is known at the molecular level. The attractive and repulsive potentials between the basic units of chromatin and the nucleoplasm are influenced by a complex combination of epigenetic modifications, such as acetylation and methylation, as well as the local physicochemical environment composed of ions and pH. Additionally, architectural proteins and other factors that topologically constrain chromatin contribute to supranucleosomal chromatin structure. All of these factors influencing chromatin packing vary along the linear genomic sequence (e.g., chromatin as a piecewise heterogeneous polymer) and in 3D throughout the nucleus. Together, they potentially drive the formation of spatially separable PDs by creating areas with similar chromatin-chromatin and chromatin-nucleoplasm interactions that are further affected by physical topological constraints.

It is important to stress that it is premature to suggest whether structurally defined PDs are related to functionally defined topologically associating domains (TADs) discovered by Hi-C: regions of tens to hundreds of kilobases with frequent intradomain interactions that exhibit a hierarchical organization (*17*–*19*). Notably, however, several properties of PDs are similar to those of sub-TADs and TADs. From ChromSTEM data, we estimated the average genomic size of PDs to be ~412.41 kbp, which is within the range of typical TAD sizes (*59*). The size of the PDs was calculated under the assumption that the voxels with the highest DNA intensity in ChromSTEM represent pure, unhydrated DNA. This assumption is likely to overestimate the genomic size of PDs. A more accurate evaluation requires additional calibration experiments to link ChromSTEM image contrast to the total mass of DNA at different pixel sizes. In addition, PDs are heritable through the process of cell division. Previous Hi-C experiments have demonstrated that transcriptional inhibition significantly alters genome connectivity within and between TADs (*60*), and similarly, the structure of PDs appears to be altered by transcriptional inhibition. Finally, the genome connectivity behavior within TADs is potentially related to the 3D conformation of the chromatin chain within PDs. However, as both ChromSTEM and PWS currently lack gene specificity, further investigation employing gene-specific labeling is required to establish a potential association between PDs and TAD-related structures. This would require the development of gene-labeling methods that are not reliant on DNA denaturation, a process that disrupts the endogenous chromatin packing structure. Providing such a link between the structural (e.g., PDs) and functional (e.g., TADs) units of chromatin organization may help elucidate the functionality of packing domains, which appear to be key structures in supranucleosomal chromatin packing.

The intricacies of the relationship between physical chromatin organization and gene transcription can be revealed by the nano-ChIA platform combined with a physics-based modeling platform. First, we demonstrate that local chromatin packing scaling influences gene expression, as predicted by our CPMC model. Gene expression is a nonmonotonic function of local chromatin packing scaling; low as well as high *D* can inhibit transcriptional processes by altering the balance between reaction rate constants and molecular mobility of transcriptional reactants. Conversely, transcription processes themselves may contribute to the packing organization of chromatin. Employing nano-ChIA, we observe a partial disruption of PDs that occurs on the order of minutes upon the inhibition of transcriptional elongation. Thus, transcription could directly influence the chromatin packing conditions to which genes are exposed. Earlier reports have suggested that transcriptionally driven DNA supercoiling and phase separation might be involved in the transcriptional regulation of chromatin conformation (*56*, *58*, *61*–*63*). However, further studies are necessary to unequivocally determine the molecular mechanisms of transcription-dependent chromatin packing regulation. Overall, the results are consistent with the view of the genome as a self-organizing system where interactions between chromatin packing behavior and transcriptional processes might represent a dynamic, self-organizing process (*64*). In this case, transcriptional reactions occur predominantly on genes associated with favorable chromatin packing environments, such as the optimal chromatin packing scaling, and at the same time, transcription processes themselves help drive the formation of domains with distinct chromatin packing behavior.

Higher-order chromatin structure changes significantly throughout the cell cycle. Mitotic chromosomes lose their cell-type-specific organization and gene expression profiles, yet both are re-established upon mitotic exit (*45*, *48*). This poses the questions of whether chromatin organization can be preserved over generations of cells and in what sequence the higher-order chromatin structures are re-established. Unfortunately, nanoimaging techniques such as ChromSTEM and biochemical methods such as chromosome conformation capture can provide only snapshots of chromatin organization, as chemical fixation is involved. Notably, the live-cell, label-free PWS module in nano-ChIA is capable of dynamically tracking chromatin organization throughout the cell cycle. Utilizing PWS, we uncovered a strong correlation between the chromatin packing scaling of progeny cells, which is also correlated with that of the progenitor cell. Interestingly, for the same progenitor cells, we observed significant synchronization of the redistribution of chromatin packing immediately after cell division. This may have significant functional consequences for dividing cells. In particular, chromatin packing scaling has been shown to be directly correlated with the phenotypic plasticity of cancer cells (*27*). Thus, the ability to inherit a more transcriptionally plastic chromatin packing structure across cell division may be a critical factor in cancer progression and the propagation of chemoresistant phenotypes.

The spatiotemporal coherence of chromatin packing scaling among progenitor and progeny cells is indicative of a heritable chromatin packing structure. This raises the question of what molecular mechanisms contribute to the re-establishment of higher-order chromatin structure across cell division. Although the molecular mechanisms of packing domain formation remain to be elucidated, most of the putative determinants are potentially heritable. The expression of ion channels, which are direct regulators of the intranuclear physicochemical environment, is genetically and epigenetically conserved across cell division. In particular, dysregulated expression and function of ion channels have been associated with the propagation of cancer phenotypes (*65*). The CTCF-cohesin complex has been shown to play a crucial role in maintaining coherent, cell-type-specific, and heritable TAD boundaries (*66*). Transcriptional memory propagation occurs through mechanisms such as mitotic bookmarking (*67*). Specifically, bookmarking transcription factors remain bound to condensed chromosomes and allow gene expression to occur during mitosis, potentially helping to re-establish transcription patterns post mitosis (*45*, *67*). Additionally, both active and repressive histone modifications have been demonstrated to be preserved throughout the cell cycle (*67*). Further investigation elucidating the contribution of these potential mechanisms to the heritability of supranucleosomal chromatin organization may, in turn, provide insights into cancer cell plasticity, the development of chemoresistance, and phenotype formation and maintenance.

Despite the strengths of the nano-ChIA platform, further development is necessary to adequately respond to the open questions explored in this work. Specifically, the labeling of active RNA polymerase is only a proxy for measuring transcription processes at specific loci. Future studies could label specific genes to assess the surrounding chromatin packing conditions using PWS and concomitantly label corresponding mRNA transcripts in the cytoplasm to obtain results more directly comparable with the CPMC model predictions and observed cellular phenotype. Additionally, we assessed the effects of transcriptional inhibition on packing domains using only PWS, which provides nanoscale sensitivity but diffraction-limited localization. Thus, this technique alone is insufficient to address the question of whether packing domains themselves are disrupted by transcriptional perturbation. To thoroughly interrogate the complex, bidirectional relationship between chromatin structure and transcription and to investigate potential mechanisms of heritability, both at all length scales, requires the future development of a ChromEM/ChromSTEM-STORM colocalization platform. Finally, as nano-ChIA currently lacks genomics information, the integration of single-cell Hi-C would be necessary to understand how local changes in chromatin structure contribute to alterations in genome connectivity. Bulk Hi-C measurements provide only a statistical description of population-wide contacts. Thus, single-cell methods would be preferred to allow a direct comparison with our single-cell nanoimaging platform. In summary, the nano-ChIA platform provides direct, high-resolution imaging of 3D chromatin structure as well as real-time, live-cell chromatin packing information over multiple length scales, highlighting the importance of combining distinct nanoscale-sensitive techniques to provide a more coherent picture of chromatin structure and function.

## Materials and Methods

### Cell culture

A549 cells were cultured in Dulbecco’s Modified Eagle Medium (ThermoFisher Scientific, Waltham, MA, #11965092). BJ cells were cultured in Minimum Essential Media (ThermoFisher Scientific, Waltham, MA, #11095080). HCT 116 cells were cultured in McCoy’s 5a modified medium (ThermoFisher Scientific, Waltham, MA, #16600082). All culture media was supplemented with 10% FBS (ThermoFisher Scientific, Waltham, MA, no. 16000044) and 100 μg/mL Penicillin-Streptomycin (ThermoFisher Scientific, Waltham, MA, # 15140122). All cells were maintained and imaged at physiological conditions (5% CO2 and 37□°C) for the duration of the experiment. All cell lines were tested for mycoplasma contamination with Hoechst 33342 within the past year. Experiments were performed on cells from passage 5–20.

### ChromEM Sample Preparation

For the EM experiment, all the cells were prepared by the ChromEM staining protocol^42^. Hank’s balanced salt solution without calcium and magnesium was used to remove the medium in the cell culture. Two-step fixation using EM grade 2.5% glutaraldehyde and 2% paraformaldehyde in 0.1M sodium cacodylate buffer (EMS) was performed: 1. Fixation at room temperature for 10 minutes. 2. Continuous fixation on ice for 1 hour with a fresh fixative. The cells were kept cold from this step either on ice or on a cold stage, and the solution was chilled before use. After fixation, the cells were thoroughly rinsed by 0.1M sodium cacodylate buffer, blocked with potassium cyanide (Sigma Aldrich) blocking buffer for 15 minutes, and stained with DRAQ5 ™ (Thermo Fisher) with 0.1% saponin (Sigma Aldrich) for 10 minutes. The excessive dye was washed away using a blocking buffer. The cells were bathed in 3-3’ diaminobenzidine tetrahydrochloride (DAB) solution (Sigma Aldrich) during photobleaching.

A Nikon inverted microscope (Eclipse Ti-U with the perfect-focus system, Nikon) with Cy5 filter sets were employed for photo-bleaching while the cells were kept cold on a custom-made wet chamber with humidity and temperature control. 15 W Xenon lamp and the red filter was used as the source of epi-illumination. With 100x objective, each spot was photo-bleached for 7 min, and fresh DAB was added to the dish for every 30 minutes. After photo-bleaching, the excessive DAB was washed away by 0.1 M sodium cacodylate buffer, and the cells were stained with reduced osmium (2% osmium tetroxide and 1.5% potassium ferrocyanide, EMS) for 30 minutes on ice to further enhance contrast. Following heavy metal staining, the cells were rinsed by DI, serial ethanol dehydrated, and brought back to room temperature in 100% ethanol. The standard procedure of infiltration and embedding using Durcupan resin (EMS) was performed. The flat embedded cells were cured at 60°C for 48 hours. The precision of the staining was tested for the entire photo-bleached cells and partially photo-bleached cells (Fig. S13).

Two types of sections were made using an ultramicrotome (UC7, Leica). For the tomography, 100 nm thick resin sections were cut and deposited onto a copper slot grid with carbon/formvar film (EMS). For investigating the chromatin structure difference with and without dexamethasone treatment, 50 nm thick resin sections were made and deposited onto copper 200 mesh grid with carbon/formvar film (EMS). The grids were plasma-cleaned by a plasma cleaner (Easi-Glow, TED PELLA) prior to use. No post staining was performed, but 10 nm colloidal gold particles were added to the 100 nm thick samples on both sides as fiducial markers for the tomography.

### Electron microscopy imaging and tomography reconstruction

A 200 kV STEM (HD2300, HITACHI) was employed for tomography data collection. High angle annular dark-field (HAADF) imaging contrast was used in the tilt series. In order to reduce the missing wedge, tilting series from – 60° to 60° on two perpendicular axes were recorded manually, with 2° step size. The pixel dwell time was kept small (~5 μs) to prevent severe beam damage during imaging. For the thin sections, a TEM (HT7700, HITACHI) was operated at 80 kV in bright field to capture high contrast chromatin data.

For the STEM HAADF tilt series, the images were aligned using IMOD with fiducial markers(*68*). 40 iterations of a penalized maximum likelihood (PML) algorithm with nonnegativity constraints in TomoPy (*69*) was employed for tomography reconstruction for each axis. The two reconstructed tomogram sets were re-combined in IMOD to further suppress the artifacts introduced by the missing cone. A nominal voxel size of 2.9 nm was used in the tomography to resolve individual nucleosomes. The 3D volume rendering was conducted using Volume Viewer in FIJI (*70*). The DNA density was used to generate color-coded nucleosome configurations, with green color dictates the lowest density, and red dictates the highest density. The chromatin binary masks were employed to generate the surface of supranuclesomal structures. The videos of example tomography and volume rendering can be found in SI Video 4, 5.

### Self-attracting polymer simulations

Coarse-grained polymer simulations were performed using LAMMPS molecular dynamics software. Chains of identical monomers were simulated using Brownian Dynamics (BD) with a Langevin thermostat. Each monomer represents a nucleosome core and has mass *m* = 1. Polymers up to ~150 kbp (1,000 monomers) were simulated. Intermonomer bonds were formed between successive monomers using the finitely extensible nonlinear elastic (FENE) potential:

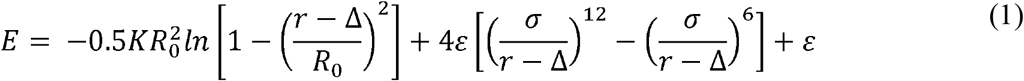

for *K* = 30.0, *R*_0_ = 1.5, *ε* = 1.0, *σ* = 1.0 and Δ = 4.0. A Lennard Jones (LJ) potential was employed to model pairwise interactions between all monomers and reinforce excluded volume effects:

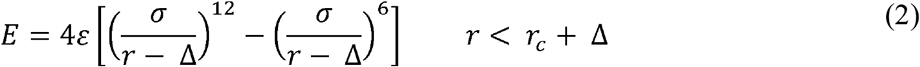

for *σ* = 1.0, Δ = 4.0, and *r_c_* = *σ* + 1.12246. *D* and *s* were modulated by tuning *ε_LJ_*, the depth of the attractive LJ well potential, between 0 to 2.5. All simulations were first equilibrated, and all mass scaling and contact probability scaling calculations were performed on trajectory files generated by subsequent production runs. Mass scaling was calculated by first counting the number of monomers within a sphere of increasing radius with an origin at the center of mass of the polymer chain and then fitting the resulting relationship between the radius of sphere and mass using linear regression. Two beads were in contact if their coordinates were within a critical distance *r_crit_* of each other in 3D space. Contact probability was calculated by summing up all observed contacts between monomers of a certain distance apart in the 1D linear chain overall 4D trajectories. A linear regression fit to the contact probability decay curve plotted against genomic distance was used to calculate contact probability scaling.

### Self-Returning Random Walk (SRRW) Simulations

The SRRW model^20^ was used to describe chromatin folding in a coarse-grained manner. The SRRW model employs steps with a continuous spectrum of step sizes. Each of these steps corresponds to about 2kb of DNA (about 10 nucleosomes), that represent the conformational freedom of a 10nm chromatin fiber, i.e. how densely or loosely packed each coarse-grained unit is. Stochastic, self-returning events are implemented through the return probability, which decays with the length of the current step size and is controlled by a chromatin folding parameter, α. We investigated the relation between contact scaling and mass scaling at varying folding states. We used the genomic range of 20kb to 200kb to probe the scaling behaviors within typical packing domains. While the contact probability as a function of genomic distance can be analyzed within such a specific window of genomic length, the mass scaling as a function of the physical radius cannot be directly mapped into the genomic window of interest. Moreover, the genomic mass that falls into a spherical probe tends to be discontinuous on the genomic sequence as the radius of the probe increases. To circumvent these problems, we used the inverse of the end-to-end distance scaling factor as an effective mass scaling factor. For a perfect mass-fractal, the end-to-end distance scaling factor is rigorously inverse to the mass scaling factor. 1000 independent conformations were generated to calculate the ensemble-averaged scaling curves. Linear regressions were used to obtain the scaling factors.

### Hi-C analysis

The Juicer analysis tool was employed to perform read alignment as well as read pairing and deduplication for each Hi-C replicates (*71*). Reads with a low mapping quality score (MAPQ < 30) were removed. Reads across replicates for each condition were merged. Juicebox was used for Hi-C contact map visualization, which was plotted with 5kb resolution. TAD sizes were calculated using the Arrowhead function from Juicer tools. All Hi-C analysis was performed on data that is publicly available through the GEO database (BJ cells: GSE81087; A549 cells: GSE92819 for control cells, GSE92811 for dexamethasone treatment). Reads from raw Hi-C data from BJ cells were mapped to the hg19 genome and Hi-C data from A549 cells was mapped to the hg38 human reference genome with MboI as the restriction enzyme. Contact probability as a function of genomic distance was calculated by normalizing observed contacts to expected contacts, (i.e., possible pairs at a given genomic distance apart). A linear regression fit on the log-log relationship between genomic distance and contact probability was performed. The mean and standard error for contact probability scaling was calculated from the slope of the regression and the standard error for this parameter estimate, respectively. In order to determine whether the difference of contact probability scaling between two treatment conditions was significant, we assumed contact probability scaling for each condition follows a normal distribution with standard deviation equal to the root-mean-square error of the regression residuals. P-values were calculated by performing a paired Student t-test assuming unequal variance.

### STORM sample preparation

Cells were grown until approximately 70% confluent on 35 mm glass-bottom Petri dishes. Cells were washed with phosphate-buffered saline (PBS) for 2 minutes then fixed with a solution of 3% Paraformaldehyde and 0.1% Glutaraldehyde in PBS for 10 minutes. Cells were washed for 5 minutes in PBS, then quenched in 0.1% sodium borohydride in PBS for 7 minutes. Cells were washed 3 times in PBS for 5 minutes each, then permeabilized in blocking buffer (0.2% Triton X-100 and 3% Bovine serum albumin (BSA) in PBS) for 20 minutes. The primary antibody (anti-RNA polymerase II, Abcam) was added to the blocking buffer to a concentration of 2.5 μg/mL and incubated for 2 hours. Cells were then washed in washing buffer (0.1% Triton X-100 and 0.2% BSA in PBS) 3 times for 5 minutes. Cells were then incubated with the secondary antibody (Alexa Fluor 647, Thermo Fisher Scientific) at a concentration of 2.5 μg/mL in blocking buffer for 40 minutes. Cells were then washed two times in PBS for 5 minutes each. Cells were imaged in standard imaging buffer with an oxygen scavenging system containing 0.5 mg/mL glucose oxidase (Sigma-Aldrich), 40 μg/mL catalase (Roche or Sigma-Aldrich) and 100 mg/mL glucose in TN buffer (50 mM Tris (pH 8.0) and 10 mM NaCl).

### STORM imaging

The STORM optical instrument was built on a commercial inverted microscope base (Eclipse Ti-U with the perfect-focus system, Nikon). The microscope is coupled to two imaging modalities. For STORM imaging, a 637 nm laser (Obis, Coherent) is collimated through a 100X 1.49 NA objective (SR APO TIRF, Nikon) with an average power at the sample of 3 to 10 kW per cm^3^. Images were collected via a 100X objectives and sent to an EMCCD (iXon Ultra 888, Andor). At least 8000 frames with a 20 ms acquisition time were collected from each sample. For PWS imaging, samples were illuminated with low NA light (0.5) and images are collected using the same 100X objective and sent through a liquid crystal tunable filter (LCTF, CRI VariSpec) and then to an sCMOS camera (ORCA Flash 4.0, Hamamatsu). The LCTF allows for spectrally resolved imaging. Images are collected between 500 nm and 700 nm with 2 nm intervals.

### PWS sample preparation

Before imaging, cells were cultured in 35□mm glass-bottom Petri dishes until approximately 70% confluent. All cells were given at least 24□hours to re-adhere before treatment (for treated cells) and imaging. A549 and BJ cells treated with Dexamethasone (Sigma-Aldrich, St. Louis, MO, D6645) were treated with a dose of 100 nM.

### PWS imaging

The PWS optical instrument is built on a commercial inverted microscope (Leica, Buffalo Grove, IL, DMIRB) using a Hamamatsu Image-EM CCD camera C9100-13 coupled to a liquid crystal tunable filter (CRi Woburn, MA, LCTF) to do hyperspectral imaging. Spectrally resolved Images are collected between 500 nm and 700 nm with 2 nm steps. Broadband illumination is provided by an Xcite-120 LED Lamp (Excelitas, Waltham, MA). For live-cell measurements, cells were imaged live and maintained at physiological conditions (5% CO2 and 37□°C) via a stage top incubator (In Vivo Scientific, Salem, SC, Stage Top Systems). As described in the Results, PWS measures the spectral standard deviation of internally optical scattering originating from nuclear chromatin, which is related to variations in the refractive index distribution (∑). Those variations in the refractive index distribution are characterized by the mass scaling or chromatin packing scaling, *D*. Therefore, D was calculated from maps of ∑. A detailed description of the relationship between ∑ and *D* is provided in the SI (*27*, *31*).

### Packing domain analysis using ChromSTEM tomography

We generated binary masks for chromatin from the ChromSTEM tomograms based on the automatic thresholding in Fiji (Otsu’s method) as reported previously (*12*). As the tomography data was obtained through STEM HAADF imaging mode, we fine-tuned the imaging processing parameters. Manually segmented chromatin masks were compared to a series of automated masks with different parameters (Fig. S14). For all chromatin masks used in this work, the following procedure was performed. First, the local contrast of the tomograms was enhanced by CLAHE, with a block size of 120 pixels. The, Ostu’s segmentation algorithm with automatic threshold was employed. Finally, we removed both dark and bright outliers using a threshold of 50 and a radius of 2 to refine the chromatin mask. All imaging processing was performed in FIJI (*72*).

On the binary chromatin masks, mass scaling analysis was performed to unveil the chromatin packing structure. The mass scaling relation M(r) is the mass of chromatin (M) contained within a sphere of radius r, and it dictates the relationship between the physical size and the genomic size of the chromatin. For a fractal structure, the mass scaling follows a power-law relation, and the scaling exponent is the packing scaling D_*MS*_. To calculate the mass scaling curve from ChromSTEM data, the total chromatin *M* (*r*) was calculated within concentric circles for each radius *r*. 100 non-zero pixels were randomly chosen on each slice of the tomography data as the origin of the concentric circles. The average MS curve was calculated from individual MS curves to reduce noise.

In addition, average MSs within 3D moving windows were used to calculate the spatial distribution of packing scaling Ds for the entire field of view. Average 2D mass scaling curve was calculated over multiple individual mass scaling curves centered on non-zero pixels located in the center region (~ 15 nm^3^) in each window. To calculate D, we employed linear regression on the average MS curve in the log-log scale, fitting from ~10 nm to ~ 30 nm. We then assigned this value to the center pixel of the 3D window to map the spatial distribution of *D*.

A contrast enhancement (CLAHE plugin in FIJI) and a flooding algorithm (MATLAB) was implemented in to segment individual PDs with similar packing scaling. We defined the boundary of each PD as the spatial separation where the mass scaling curve deviates from a fractal behavior, and the distance from the center of the PD to the boundary is the domain radius R_*f*_. One of the three criteria has to be met if the mass scaling curve deviates from a fractal behavior: 1. The linear fit of the power law from 11.7 nm to 33.3 nm is 5% different from mass scaling curve (bad fitting); 2. The slope of the mass scaling curve reaches 3 (space filling). 3. The curvature (second derivative) of the mass scaling curve reaches 2 (non-linaer). If all criteria are satisfied in one mass scaling curve, we chose the smallest value to be R_*f*_. An example of such process can be found in Fig. S15. We calculated the average mass scaling from individual mass scaling centered on all the voxels within one packing domain, and quantified the D and R_*f*_. Assuming the highest intensity in the tomograms represents 100% unhydrated DNA (density = 2g/cm^3^), and the average molecular weight for nucleotide is 325 Da, we calculated the highest mass (*m*) per voxel (*dr* = 2 nm) to be ~15 bp. We further calculated the average genomic size of PDs to be 421.41 kbp by 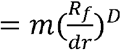, with *R_*f*_* = 106.0 nm and *D* = 2.58.

### Chromatin fractal dimension comparison for cells with dexamethasone treatment

TEM images of 50 nm thin sections were used in the analysis of chromatin packing alterations induced by the dexamethasone treatment for 32 hours. Unlike STEM HAADF imaging mode, the TEM bright-field contrast attenuates following Beer’s law,

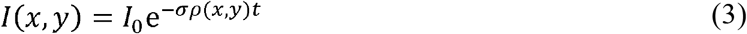

Where *I*(*x*,*y*) is the TEM image intensity distribution, *I*_0_ is the incident beam intensity, *σ* is the absorption coefficient, *ρ*(*x*,*y*) is the density distribution, and *t* is the section thickness. In our experiment, *I*_0_, *σ*, *t* were controlled to be constant for all images, only the chromatin density *ρ*(*x*,*y*) contributes to the final image intensity *I*(*x*,*y*). To obtain the density fluctuation, *ρ*_Δ_(*x*,*y*), we took the negative logarithm of all the TEM images directly and subtracted the mean value. At the same time, the incident beam intensity *I*_0_ is canceled out.

The two-dimensional autocorrelation function (ACF) was calculated using the Wiener-Khinchine relation as:

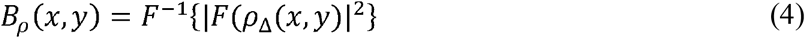

Where *F*^−1^ and *F* is the inverse Fourier, and the Fourier transforms, and the *ρ*_Δ_ is the fluctuating part of the chromatin density. To minimize the noise, a rotational average of *B_ρ_*(*x*,*y*) was taken to obtain the final form of the ACF *B_ρ_*(*r*), representing the correlation of chromatin density as a function of spatial separation *r*. Notice that mathematically, a fractal structure can be characterized by a power-law ACF, *B_ρ_*(*r*) ~ *r*^*D*–3^, with *D* being the fractal dimension. For the chromatin reconstructed by ChromSTEM, the mean ACF *B_ρ_*(*r*) was averaged over the ACFs of each virtual 2D slices and plotted in log-log scale. Linear regression was performed from 50 nm to 100 nm to obtain the slope *p*. The chromatin packing scaling *D* was calculated by 3 + *p*.

Each nucleus was carefully segmented manually in FIJI and the chromatin packing scaling *D* was calculated through the ACF analysis within the nucleus. Multiple cells in the control group (BJ: n = 31, A549: n = 8) and the treated group (BJ: n= 31, A549: n = 10) were compared.

### Chromatin Packing Macromolecular Crowding (CPMC)

As previously introduced in the CPMC model, at specific chromatin packing scaling D, the average expression of a group of genes can be approximated as the product of two components, i.e., the probability of the genes to be on the accessible surface *P_s_* and the average mRNA expression rate of these genes 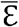:

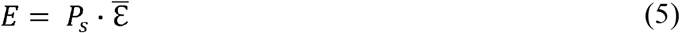

Based on the power-law mass scaling (fractal property) of the chromatin, *P_s_* is determined by the mass of fractal *M_f_*, the mass of the elementary particle in the chromatin fractal and the chromatin packing scaling D as:

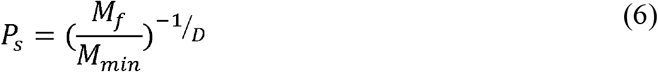

The average expression rate of mRNA for genes with specific molecular factors 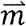 can be evaluated by integrating mRNA expression rate 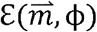 with crowding density distribution ϕ. This is performed modeling transcription as a series of chemical reactions and then solving the steady-state network of equations as described by *Matsuda et al*. This systems biology method incorporates results from Brownian Dynamics and Monte Carlo simulations to study the effects of increased crowding and molecular regulators of transcription on diffusion and binding of transcriptional reactants, respectively. If the probability distribution function of ϕ is f(ϕ), the average mRNA expression rate is, therefore:

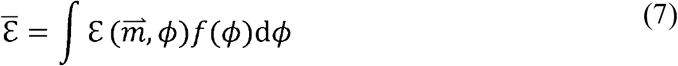

Without the loss of generality, crowding density can be assumed to follow a Gaussian distribution. Thus, *f*(*ϕ*) can be approximated as 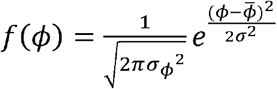, where *σ*^2^ is the variance of crowding density and 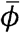 is the average crowing density of the entire nuclear. The *σ_ϕ_*^2^ is determined by D as: 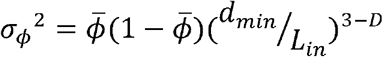, where *d_min_* is the diameter of the elementary particle in the chromatin (the nucleosome in this case) and *L_in_* is the length of the interaction volume whose nanoenvironment can affect the transcription of a single gene.

### Data analysis for multi-modal STORM-PWS studies

STORM images were reconstructed using the ThunderSTORM plugin for FIJI (*73*). Maps of chromatin packing scaling (*D*) from raw PWS images were created using a custom analysis script in MATLAB that has been described in detail in previous works (*31*) Colocalization between PWS and STORM images was achieved through alignment of wide field reflectance images collected on each separate imaging arms. Errors caused by sample drift during imaging sequences were first corrected in ThunderSTORM; any additional corrections were applied manually as needed. To create plots in Fig. 4 and Fig. S8, the average chromatin packing (*D*) is calculated from the PWS data (in red), and the average local Pol II concentration was calculated from STORM data (in green) in each pixel (130 nm X 130 nm). Data points with similar *D* are grouped together (*D* within 0.025) and plotted in Fig. 4D. The circles represent the means, and error bars are a standard error between regions.

In order to do a spatial analysis of packing domains and POL II locations, maps of D measured by PWS microscopy were first binarized using the function ‘imbinarize’ in MATLAB with adaptive thresholding. This allows the segmentation of packing domains. Next, the Euclidean distance transform of the binarized image was calculated using ‘bwdist’ to find the distance between each pixel and the nearest packing domain. Positive distances denote pixels outside of packing domains, and negative distances denote pixels inside of packing domains. Next, all pixels containing no Pol II (as visualized by STORM) were removed. The distances from pixels with POL II density higher than the mean (i.e., the most Pol II rich regions) were plotted in a violin plot (Fig. 4E).

### Halting Transcriptional Elongation

BJ cells (Fig. 3) and A549 cells (Fig. S10) were treated with 5 μg/mL of Actinomycin D (Sigma, A9415), which inhibits transcription by halting elongation of the transcribed RNA (*42*, *43*). Immediately after the introduction of Actinomycin D, PWS images were collected continuously for 10 minutes (1 image collected every ~15 seconds). All cells within each field of view were analyzed. Average nuclear *D* was tracked. Additionally, maps of *D* from PWS were thresholded and segmented to find the PD projection fraction. The PD projection fraction indicates the fraction of the 2D projection of the nucleus, measured by PWS, occupied by PDs. A PD projection fraction of 1, would indicate the entire nucleus is filled with PDs. Changes in the PD projection fraction were observed over time.

### Characterizing Heritability of Chromatin Packing with PWS

HCT 116 cells were monitored with PWS for a period of 24 hours. Images were captured every 15 minutes. Ten cells which could be observed dividing during the 24-hour period were identified and chosen for analysis. *D* was tracked for 3 hours prior to cell division and at least 6 hours after cell division.

To analyze spatial correlations between progeny cells and their progenitors, histograms of *D* at all pixels within the nucleus were compared. First, each cell was analyzed at each time-point. A histogram of *D* was calculated with 10 evenly spaced bins with widths of 0.15. Each histogram was analyzed by being normalized by the average histogram of all cells at the same time point. For example, a histogram of cell #1, three hours before cell division, was normalized by the average of all cells (N = 10) histograms three hours before cell division. Therefore, these normalized histogram ratios focused on each cell’s specific deviation from the mean of the population at a specific time-point. See Fig. S12 for a step by step explanation for how histograms were calculated and normalized. The Pearson correlation coefficient was calculated in MATLAB with the ‘corrcoef’ function comparing every pair of progeny cells at each timepoint. Also, all progeny cells at three hours post-division were compared to all progenitor cells at three hours pre-division. Three hours was chosen in order to compare cells pre- and post-division at relatively stable time points (i.e., not during cell detachment or nuclear splitting).

To analyze temporal correlations in averaged nuclear *D*, the Pearson correlation coefficient was calculated in MATLAB with the ‘corrcoef’ function to compare time-dependent changes in D between each of the progeny cells (after division). To compare progeny cells to the progenitor cells, 2 hours was compared between all progeny cells and progenitors. The two-hour time period was 1 hour pre- (for progenitors) and post- (for progeny cells) division. A 1-hour “buffer” around the time of cell division was chosen since the measured *D* was highly dynamic during this time and subject to artifacts since the cell was in the process of partially detaching from the glass substrate.

## Acknowledgments

The authors would like to thank Dr. Mark Ellisman for advice and sharing unpublished results. This research was supported in part through the computational resources and staff contributions provided by the Genomics Compute Cluster which is jointly supported by the Feinberg School of Medicine, the Center for Genetic Medicine, and Feinberg’s Department of Biochemistry and Molecular Genetics, the Office of the Provost, the Office for Research, and Northwestern Information Technology. The Genomics Compute Cluster is part of Quest, Northwestern University’s high-performance computing facility, with the purpose to advance research in genomics. An award of computer time was provided by the INCITE program. This research used resources of the Argonne Leadership Computing Facility, which is a DOE Office of Science User Facility supported under Contract DE-AC02-06CH11357. The EM work made use of the BioCryo facility of Northwestern University’s NU*ANCE* Center, which has received support from the Soft and Hybrid Nanotechnology Experimental (SHyNE) Resource (NSF ECCS-1542205); the MRSEC program (NSF DMR-1720139) at the Materials Research Center; the International Institute for Nanotechnology (IIN); and the State of Illinois, through the IIN.

