## Supplementary material for "Nanoscale Chromatin Imaging and Analysis (nano-ChIA) platform bridges 4-D chromatin organization with molecular function": SI

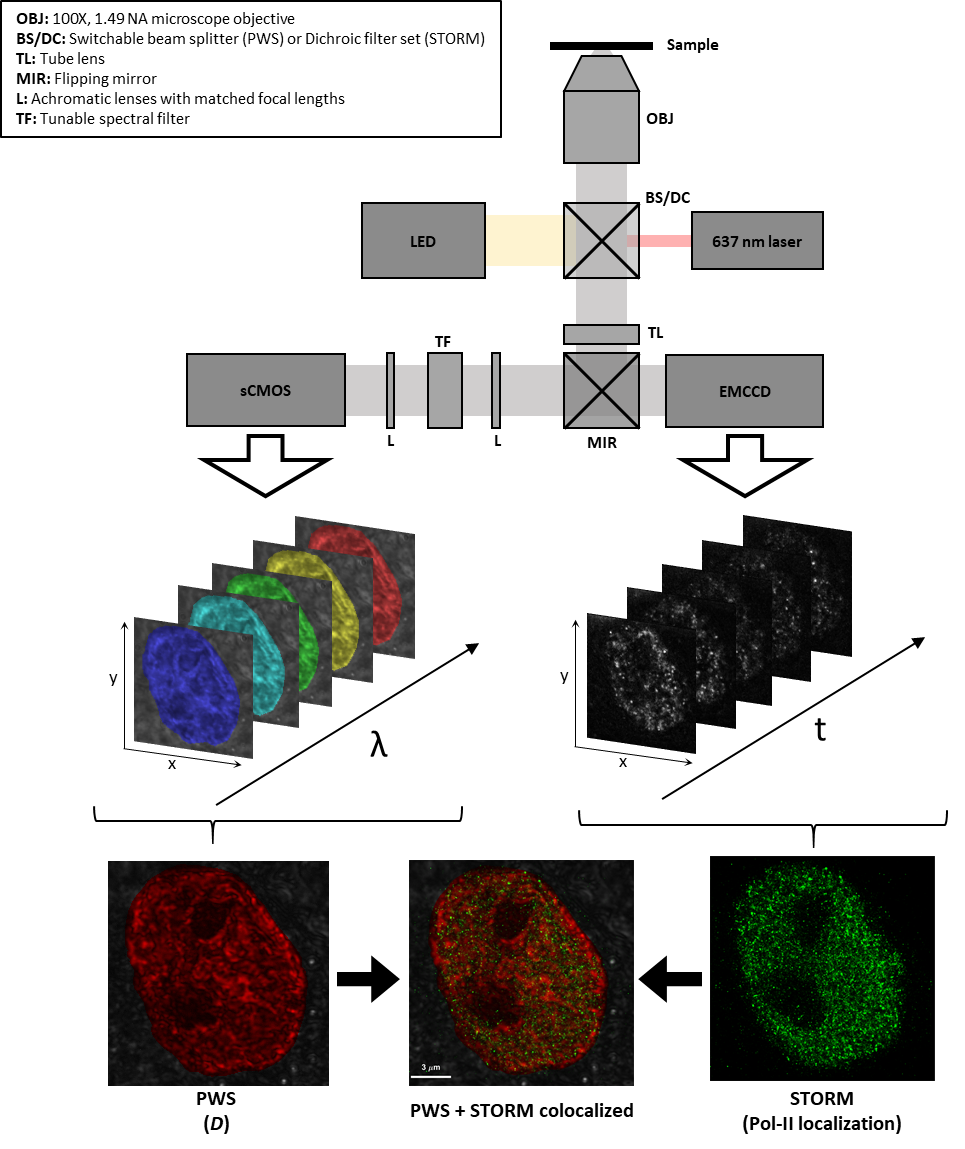

**Fig. S1. Optical schematic of the multi-modal STORM-PWS optical microscope.** During PWS acquisition, broadband light from an LED is introduced onto the sample via a 50/50 beam splitter. The light is collected and sent to an EMCCD or sCMOS camera though a tunable spectral filter. During STORM acquisition, 637 nm monochromatic red laser light is introduced onto the sample via a dichroic filter set. Fluorescence light is collected and sent to an EMCCD camera. For PWS, multiple spectrally resolved images are acquired and analyzed to create a map of *D*. For STORM, multiple frames of fluorescence emission events are acquired and analyzed to reconstruct a super-resolution fluorescence image of labeled Pol-II. The reconstructed STORM image and calculated *D* map are finally combined into a single image.

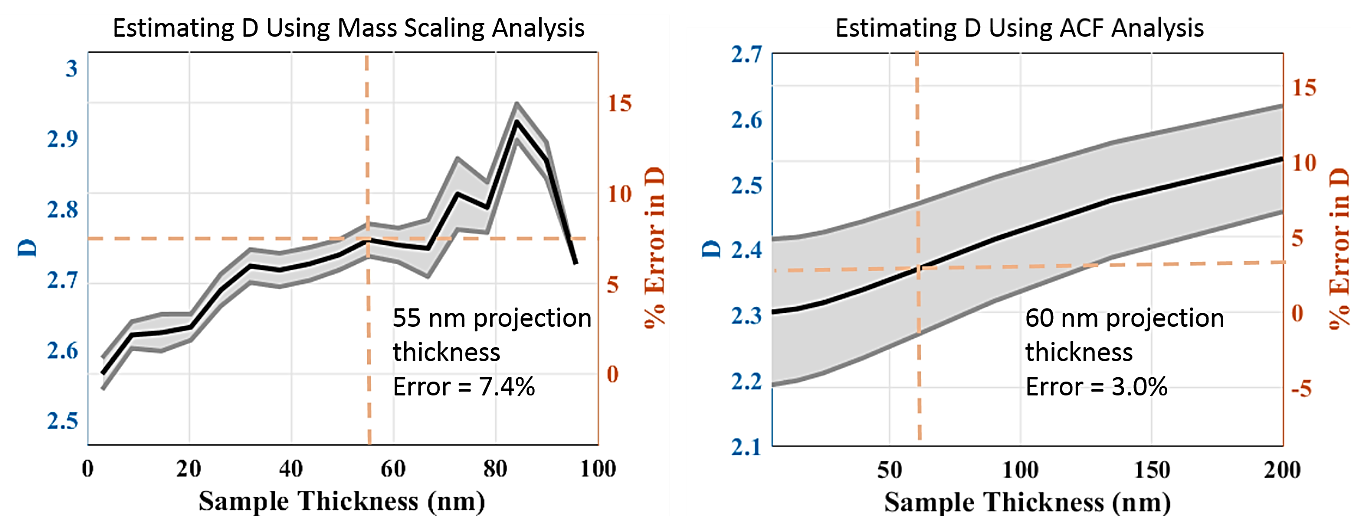

**Fig. S2. Estimating D from the projection of chromatin at varying thicknesses.** For the mass scaling analysis, we created a series of projection images with incremental thickness by projecting the virtual 2D slices of the 3D tomograms of A549 chromatin masks, respectively. Mass scaling curves were calculated first. Then linear regression was performed within 11.6 nm to 50 nm. For the ACF analysis, a random media was generated with 600^3 voxels, 5nm pixel size along each dimension and ACF characterized by chromatin with a fatal dimension of 2.3. The generated medium is 4 um in each dimension to ensure the fractal range could be fit given the numerical limitations. Then, 2D projects of the 3D volume were averaged, and the slope of their ACF was fit to a power law. All fits were done within the same range from 50-100nm. For both cases, the average *D* for each thickness is plotted. For mass scaling analysis, the estimated D is 7.4% larger than the ground truth for a 55 nm thick projection. For ACF analysis, only a 3% error in D will occur for a 60 nm thick projection image of chromatin.

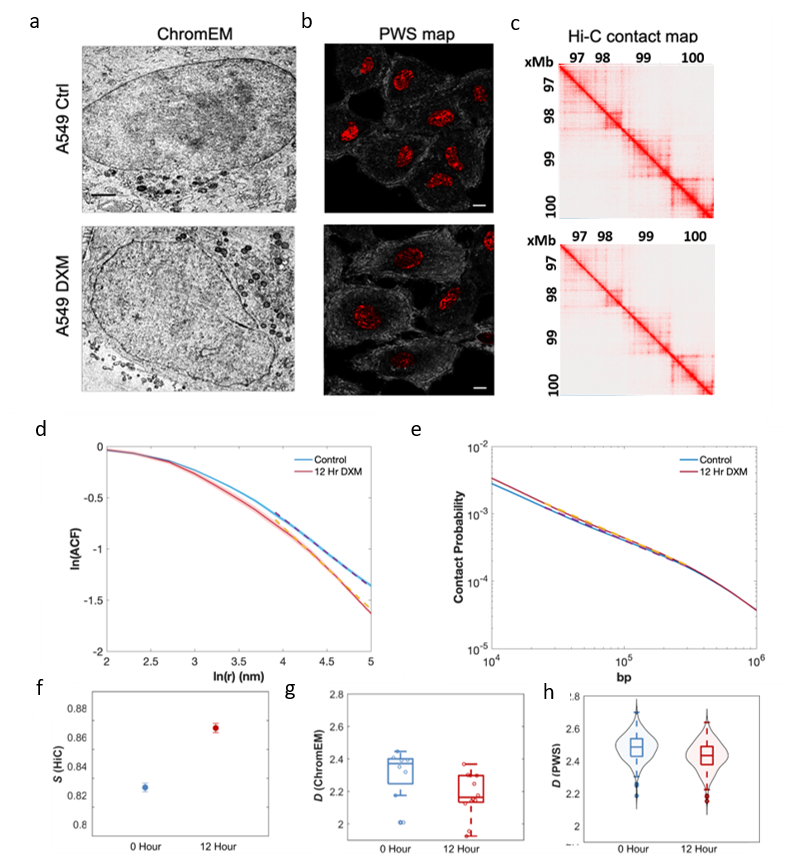

**Fig. S3. Measuring chromatin packing scaling alterations induced by dexamethasone (DXM) in A549 cells.** (a-b) nano-ChIA platform characterization of A549 chromatin with and without DXM treatment. From left to right: TEM images of chromatin structure with ChromEM staining, scale bar: 1µm. PWS map of chromatin packing scaling, scale bar: 10 µm. Qualitatively, both ChromTEM and PWS images show that DXM treatment makes chromatin packing more homogenous. (c) Hi-C contact map of human chromosome 1 rendered with 5kb resolution. (d) ACF analysis using ChromTEM on A549 cells. The average ACF of the control group (blue) is significantly different from the average ACF of the treated group (red). The shaded regions represent standard errors. *D* was measured inside the first fractal domain (50 nm to 100nm) by a linear regression fit of the ACF in the log-log scale. (e) Contact probability analysis performed on whole genome intrachromosomal Hi-C contact data. Contact probability scaling (*s*) was calculated from a linear regression fit (dotted line) of the contact probability curve in the log-log scale between genomic distance 10^4.4^ and 10^5.5^ bp. (f–h) Chromatin packing scaling alterations induced by DXM treatment measured using ACF analysis of TEM images and PWS and changes in contact probability scaling of Hi-C contact data. Across the platform, consistent changes were observed in chromatin packing scaling upon treatment.

**Fig. S4. Analyzing differences in contact probability scaling upon dexamethasone (DXM) treatment.** Comparing distributions of contact probability scaling for (a) A549 and (b) BJ cells calculated from Hi-C contact matrices. Assume the linear regression fit used to calculate contact probability scaling follows a normal distribution $\mathcal{N(}\mu_{s},\sigma_{s})$ where mean contact probability scaling, $\mu_{s}$, is the slope of the regression and standard deviation, $\sigma_{s}$is the root-mean-square error (RMSE) of the residuals. (a) Contact probability scaling is significantly different between control, and 12-hour DXM treated A549 cells (P = 0). (b) Hi-C analysis from BJ cells also demonstrates a significant difference between control and 16-hour DXM treated cells (P = 1.98E-115) as well as between control and 32-hour DXM treated cells (P = 0).

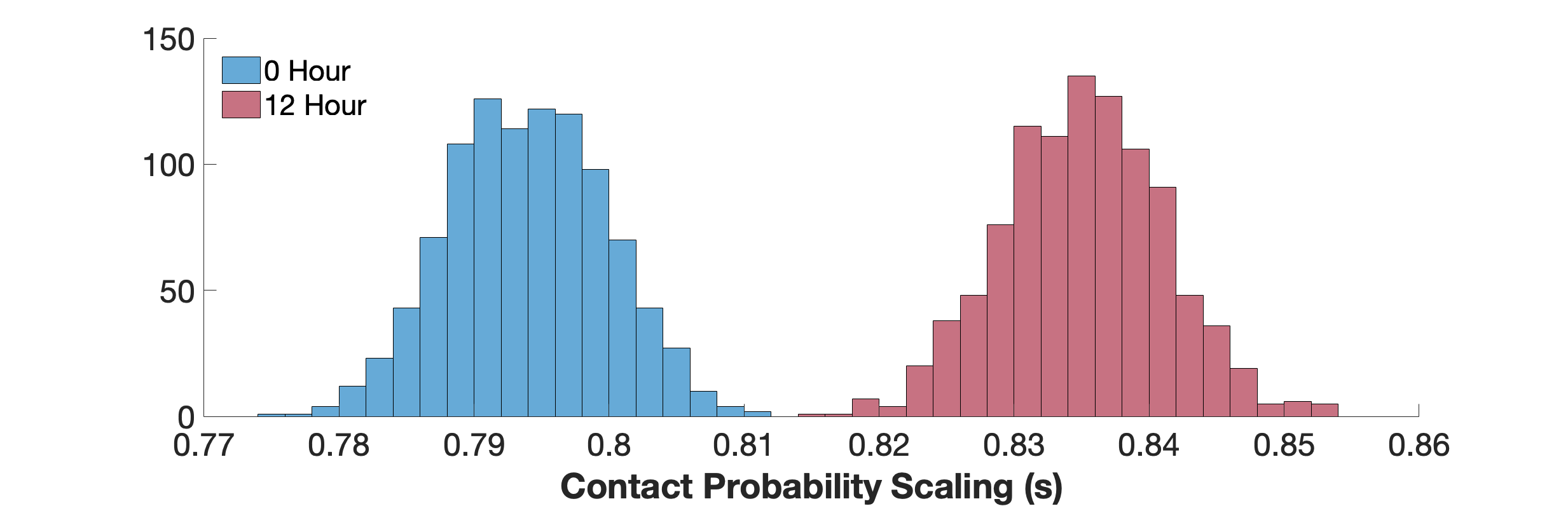

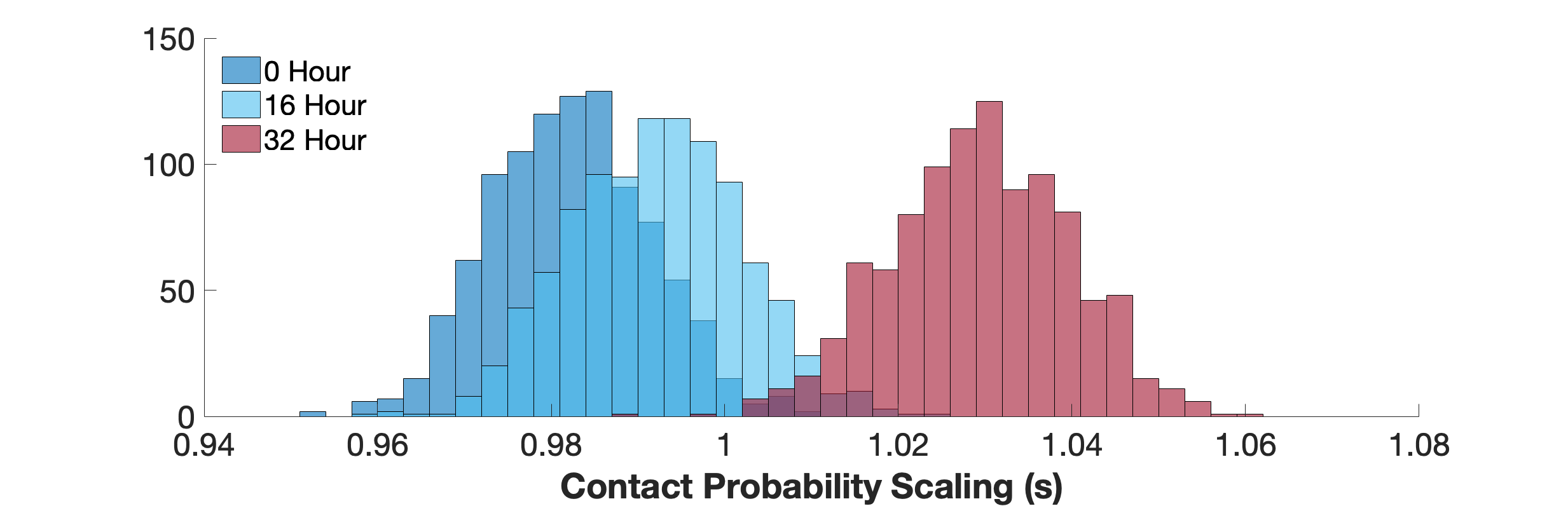

**a**

**b**

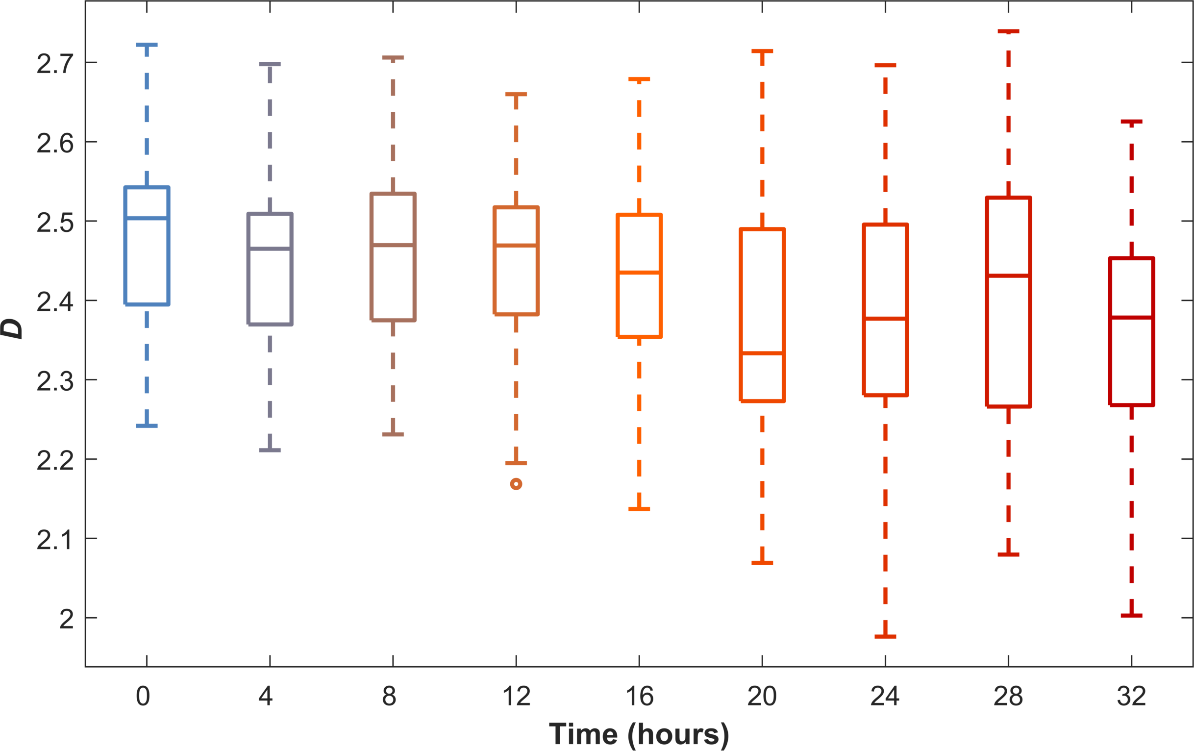

**Fig. S5. Measuring chromatin packing scaling alterations induced by dexamethasone (DXM) in BJ cells.** A subset of cells shown in Fig. S3 was measured every 4 hours during DXM treatment to demonstrate the live-cell, time-resolved measurement abilities of PWS microscopy. Here, 81 fields of view were repeatedly imaged with PWS every 4 hours. Between 67 and 96 cells were measured at each time point. Starting at 12 hours, each time point showed a statistically significant change in *D* (P < 0.05) from the 0-hour time point.

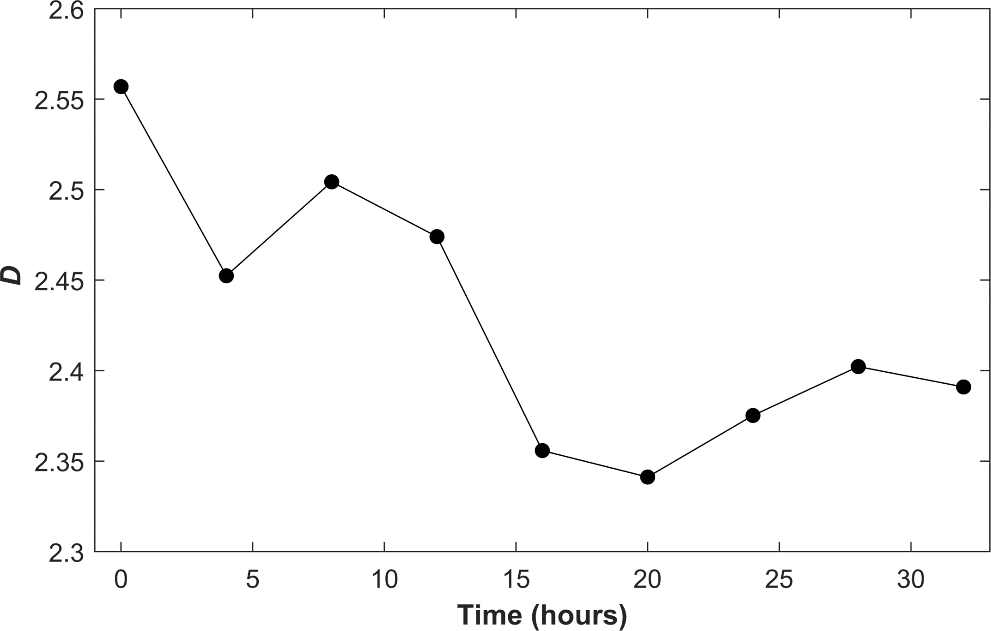

**Fig. S6. Single-cell analysis of chromatin packing scaling alterations induced by dexamethasone (DXM) in a BJ cell.** Average *D* from a single BJ cell treated with DXM from the population shown in Fig. S3 is plotted against time, demonstrated the ability of PWS microscopy to perform time-resolved, single-cell analysis of chromatin packing scaling *D*.

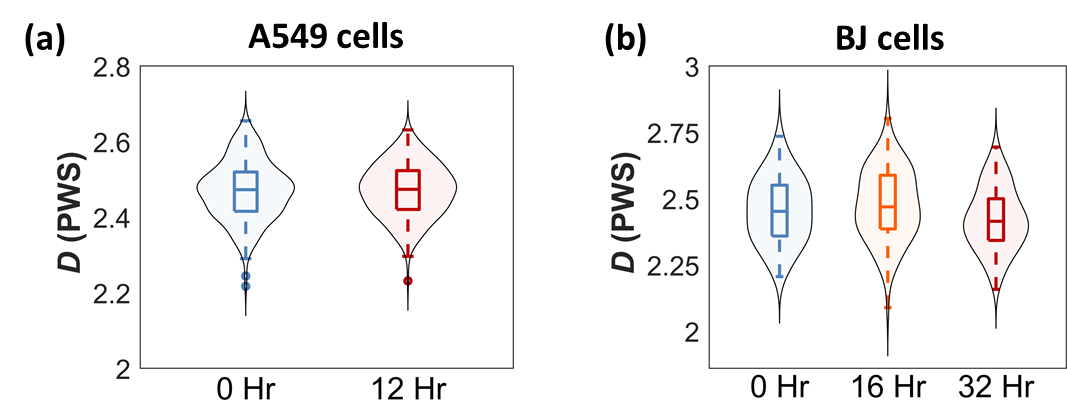

**Fig. S7 Measuring chromatin packing scaling alterations over time in untreated in A549 and BJ cells.** (a) Chromatin packing scaling *D* measured in untreated live A549 cancer cells to act as a control experiment to DXM treated cells shown in Fig. 3i. There is no observable change in *D* over 12 hours. (b) Chromatin packing scaling *D* measured in untreated live BJ fibroblast cells to act as a control experiment to DXM treated cells shown in Fig. S3d. There is no observable change in *D* over 16 hours, and while there appears to be a small decrease in *D* after 32 hours, the change is non-significant (P > 0.05).

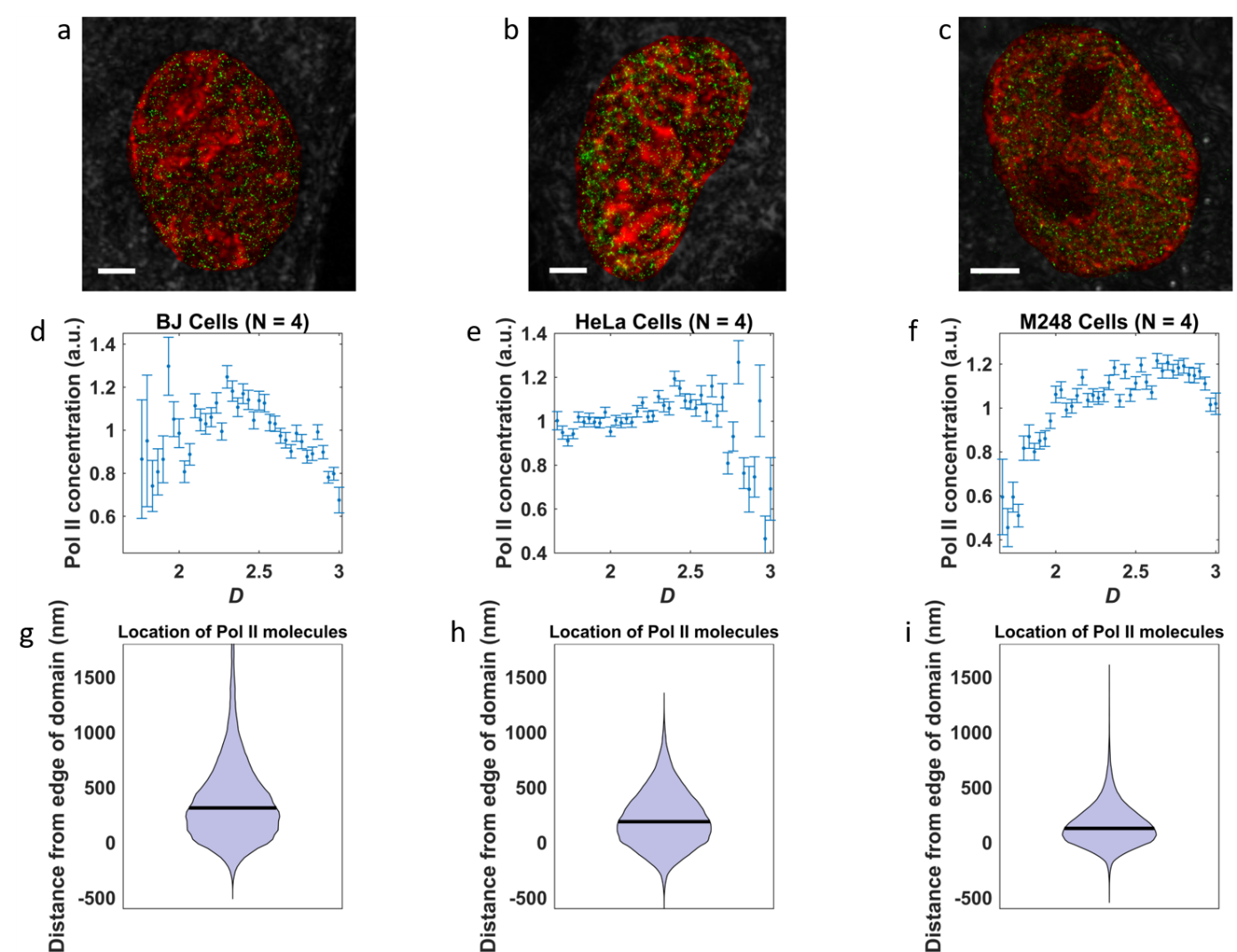

**Fig. S8. Quantifying chromatin structure and transcription using the nano-ChIA platform.** (a-c) Representative images from differentiating BJ fibroblast cells, HeLa cells and A2780.M248 (M248) ovarian cancer cells. Red shows chromatin packing scaling measured by PWS, and green shows the locations of Pol-II molecules visualized by STORM. Scale bar: 3$\mu m$. (d-f) The relationship between Pol-II density and local *D* is plotted for all three cell lines. While there is considerable variation from cell-type to cell-type, the nonmonotonic correlation is consistent. (g-i) Violin plots show the distribution of distances between highly enriched Pol-II regions and their nearest PD. In all three cell lines, it appears Pol-II tends to group near the border of PDs.

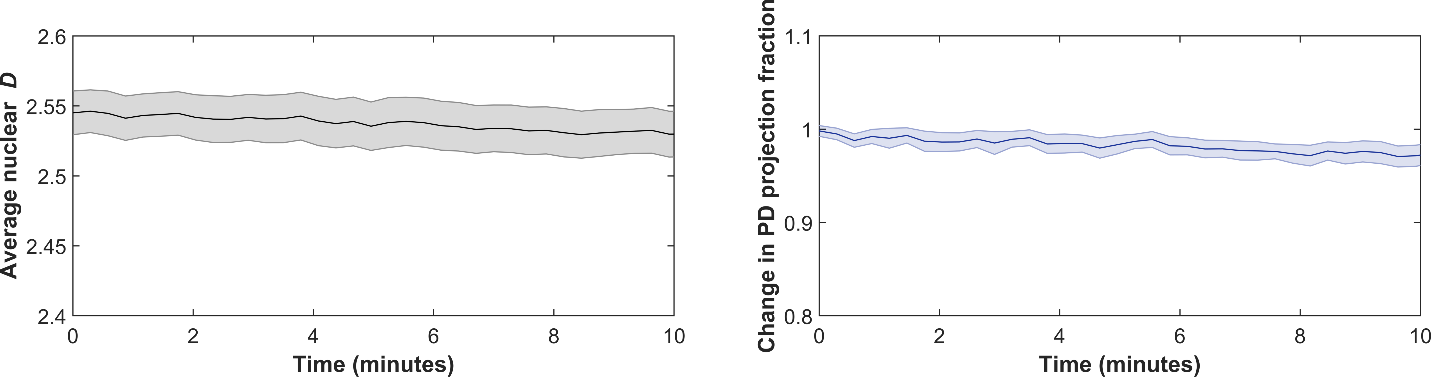

**Fig. S9. Quantifying changes in chromatin structure without perturbation.** Untreated BJ cells were monitored continuously with PWS for ten minutes to act as a control experiment for **Act-D** treatment (a) There is no noticeable change to average nuclear *D* over ten minutes. (b) There is no change to the PD projection fraction over ten minutes.

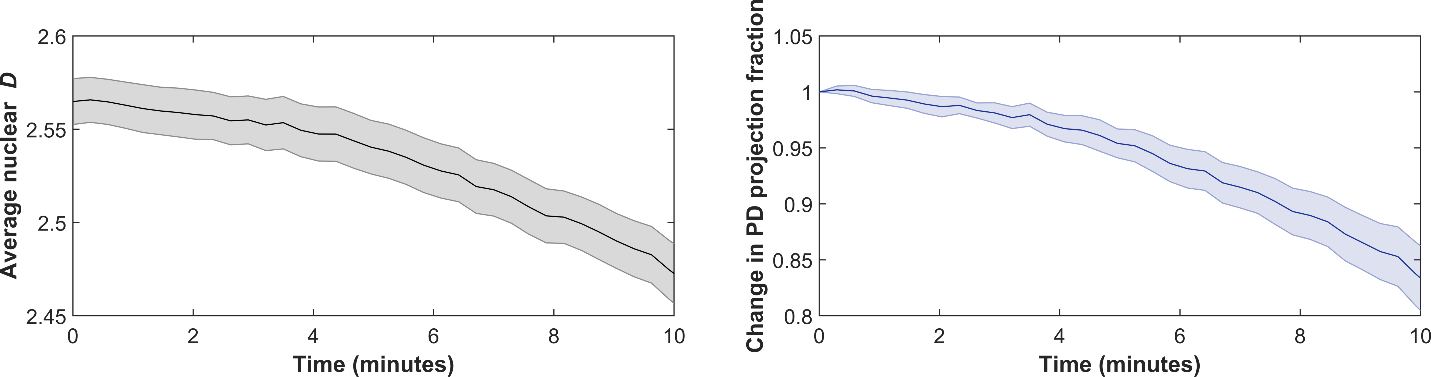

**Fig. S10. Quantifying changes in chromatin structure upon treatment of A549 cells with Actinomycin D.** A549 cells were monitored continuously with PWS for ten minutes after treatment with Act-D. (a) Average nuclear *D* drops significantly over ten minutes. (b) There is a significant drop in the PD projection fraction.

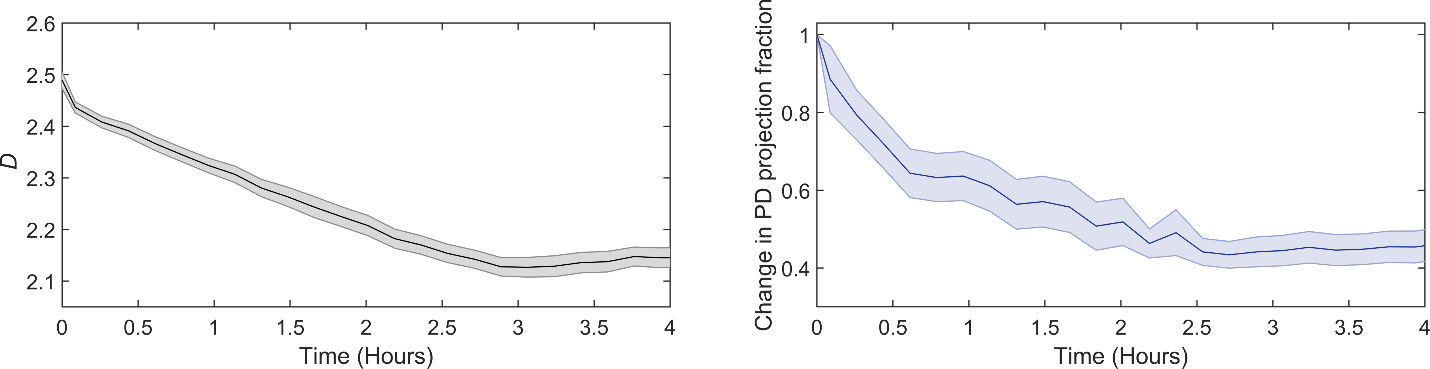

**Fig. S11. Quantifying changes in chromatin structure upon treatment of BJ cells with Actinomycin D over long timescales.** BJ cells were monitored continuously with PWS for four hours after treatment with Act-D. (a) Average nuclear *D* drops significantly over the first three hours and then appears to plateau. (b) There is a significant drop in the PD projection fraction, however the PD projection fraction never drops below ~40% its initial value. It should be noted that since, Act-D inhibits all transcription, these cells eventually go through apoptosis.

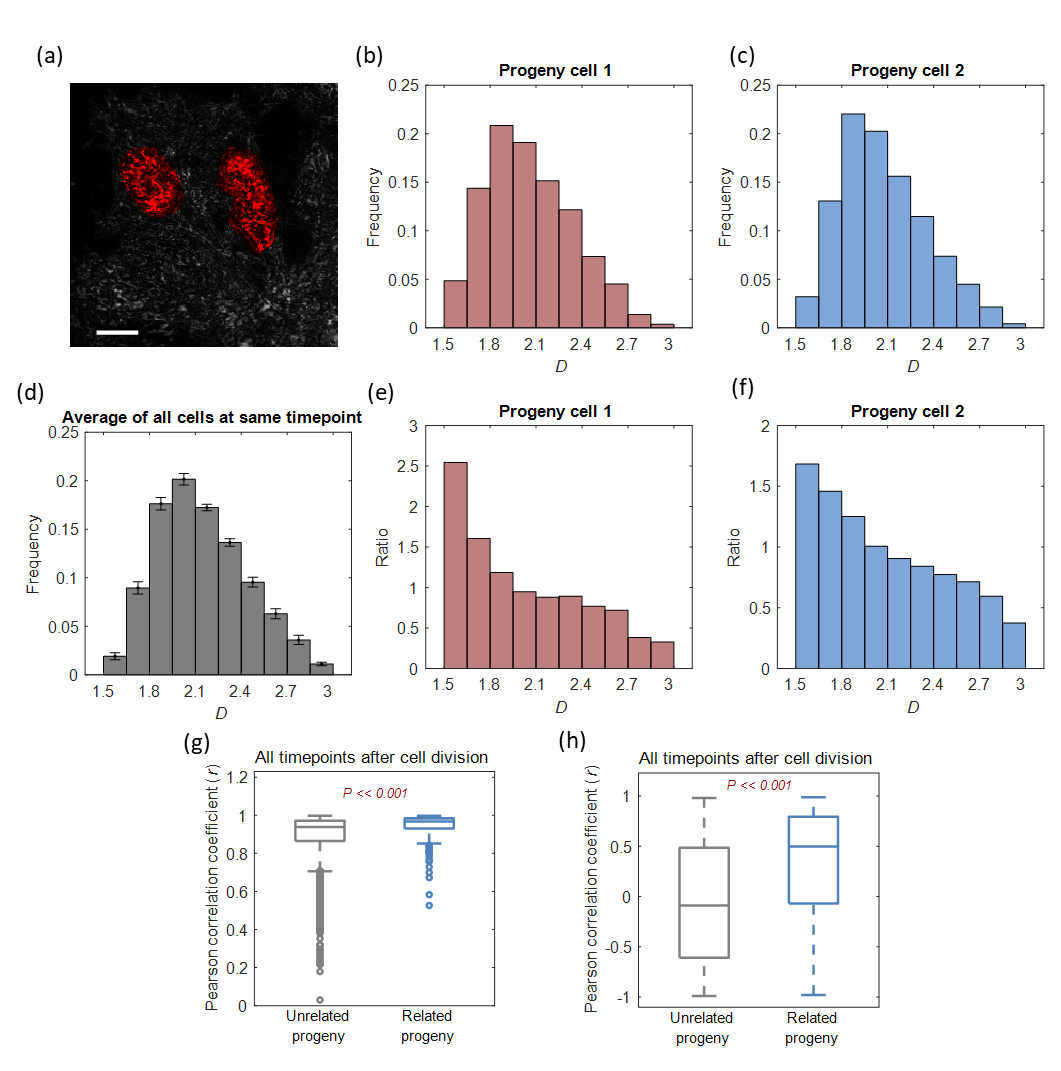

**Fig. S12. Histogram calculations for heritability analysis.** (a) A PWS image of two HCT 116 cells originating from the same progenitor cell, imaged 5 hours after cell division, was completed. (b & c) A histogram of *D* values from the two HCT116 cells shown in (a). (d) The average histogram for all cells 5 hours after cell division is shown. (e & f) The ratio of histograms shown in (b) and (c) and the average histogram is shown in (d). This ratio of histograms shows how these two cells’ characteristics deviate from the population mean. (g) A boxplot is showing the Pearson correlation coefficient calculated from the histogram of all cells at all time points after cell division. While cells originating from the same progenitor have significantly higher correlation, even unrelated cells have a high correlation (>0.5) since all cells are of the same type. (h) A boxplot is showing the Pearson correlation coefficient calculated from the ratio-histogram of all cells at all time points after cell division. Since each cell’s histogram is divided by the mean, unrelated cells tend to not correlate, while related progeny cells are significantly more correlated.

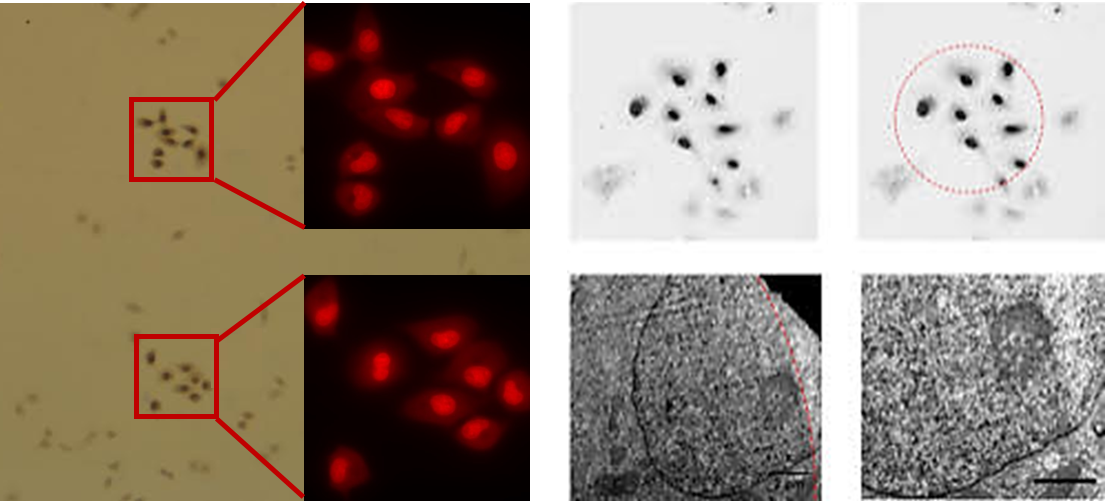

**Fig. S13. ChromEM staining reveals DNA distribution inside the A549 cell nuclei.** Draq5^TM^ was used to label the double-strand DNA inside the nucleus. Upon excitation, the Draq5 molecule will release single oxygen, which reacts with the diaminobenzidine-tetrahydrochloride (DAB) solutes to form electron-dense precipitates. The DAB precipitates are further enhanced by reduced osmium to create contrast for EM imaging. As shown in the fluorescent images of A549 labeled by Draq5 and bright-field optical images of the resin-embedded cells, only cells with photo-oxidation show darker contrast. The staining area has high precision, as indicated by a partially photo-oxidized cell. Within the nucleus, the boundary of the stained area follows the shape of the light spot.

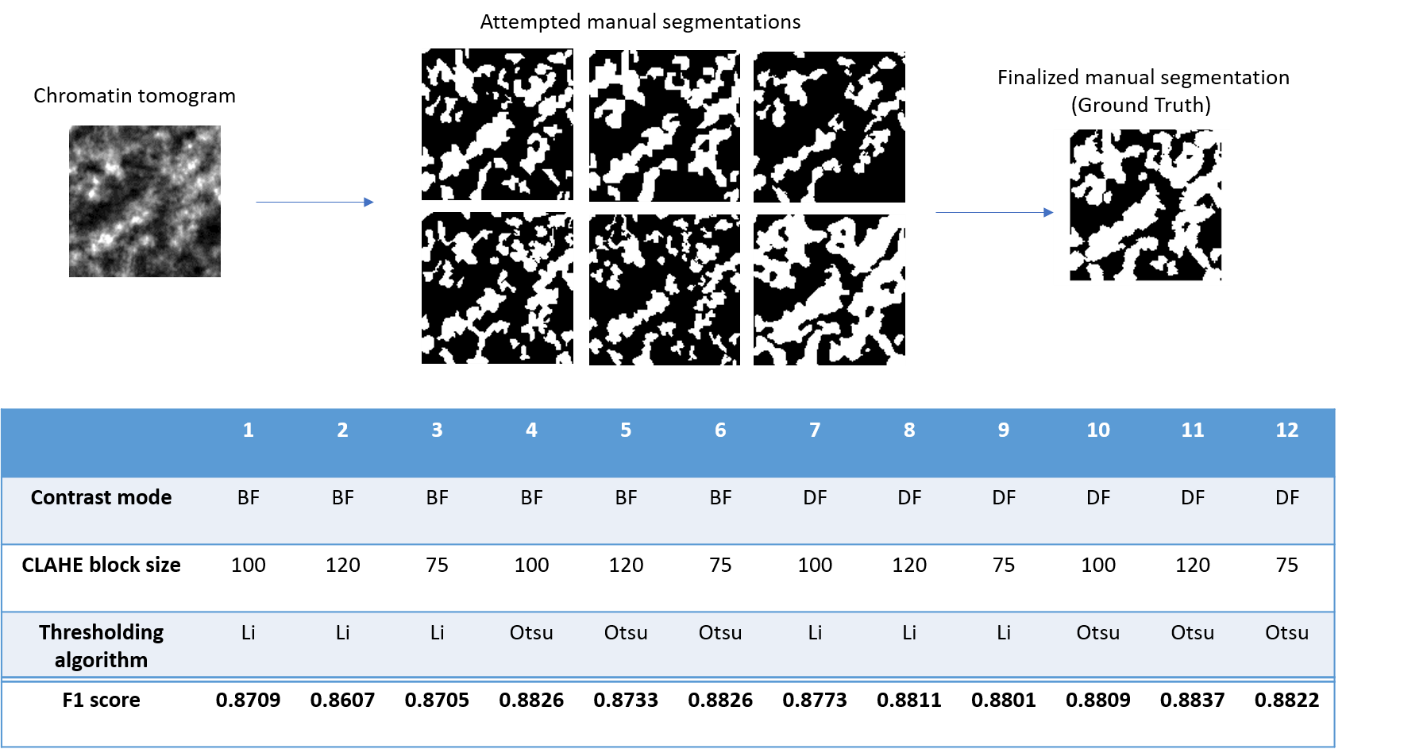

**Fig. S14. Automatic chromatin mask segmentation parameters optimization.** In the automatic chromatin mask segmentation algorithm, three parameters, contrast mode, CLAHE block size, and thresholding algorithm, need to be optimized for best segmentation accuracy quantified by F1 score. In six independent analysis, we manually segmented the chromatin mask from the same tomogram. We averaged all the masks, and pixels with values greater than 0.2 were considered chromatin, while pixels with values smaller than 0.2 were considered not chromatin. The finalized manual segmentation was used as the ground truth. Either bright field (BF) or dark field (DF) can be used in contrast mode to initiate segmentation. We chose three block size (in pixels), 75, 100, and 120, for local contrast enhancement (CLAHE), and employed either Li’s or Otsu’s thresholding algorithm in FIJI. In combination, we experimented with 12 different sets of parameters. We calculated the F1 score for each set of parameters, and determined the optimized parameters are DF as contrast mode, 120 pixels for CLAHE, and Otsu’s algorithm.

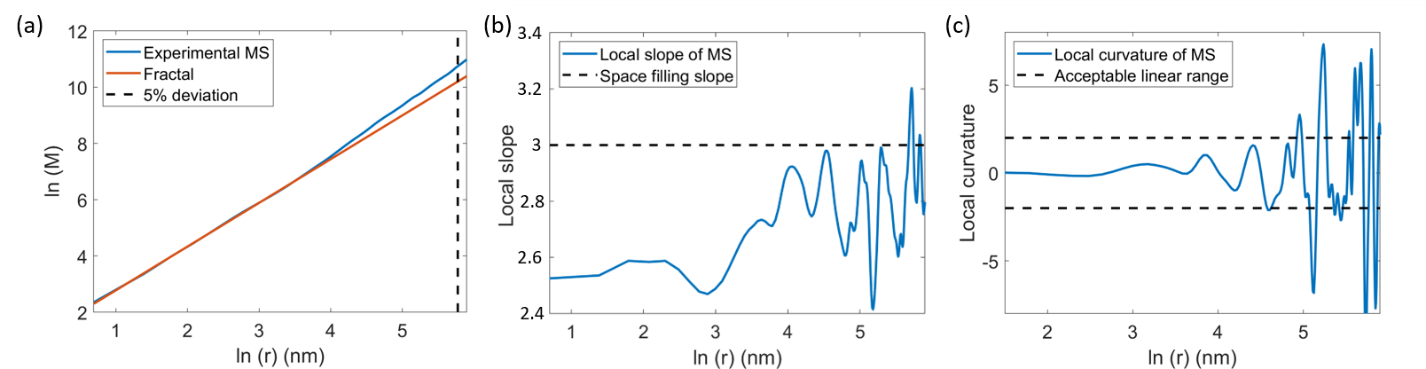

**Fig. S15. Determining the packing domain radius (*R_f_*).** Three criteria are employed in calculating *R_f_* from the mass-scaling (MS) curve for individual domains. (a) Linear regression was conducted for the mass scaling curve from 2 nm to 32 nm range. The spatial separation where the linear regression has 5% deviation from the experimental MS curve (dashed line) is defined as *R_fit_*. (b) The local slope of the experimental MS curve was estimated by moving window linear regression (5 pixels per window). The spatial separation where the local slope reaches 3 (dashed line) is defined as *R_space_filling_*. (c) The local curvature of the experimental MS curve was again calculated by moving window linear regression on the local slope curve (5 pixels per window). The spatial separation where the absolute value of the local curvature exceeds 2 (dashed line) is defined as *R_non_linear_*. *R_f_* is defined as the smallest value among *R_fit_*, *R_space_filling_*, and *R_non_linear_*. For cases where either *R_fit_*, *R_space_filling_*, *R_non_linear_* does not exist, we use infinity for that particular parameter.

**SI Video 1 – Single BJ cell treated with Dexamethasone measured with PWS for 32 hours.** The average nuclear *D* of this cell is shown in Fig. S3.

**SI video 2 – HCT 116 cancer cells measured by PWS every 15 minutes during mitosis.** Data collected from this cell is shown in Fig. 5.

**SI video 3 – HCT 116 cancer cells measured by PWS every 15 minutes during mitosis.** Data collected from this cell is shown in Fig. 5.

**SI video 4 – Tomography reconstruction of an interphasic A549 cancer cell.**

**SI video 5 – Volume rendering from tomography of the cell in SI video 4**.

| **Fixed Parameters** | **Description** | **Value** |
| --- | --- | --- |
| $\boldsymbol{K}_{\boldsymbol{D}}$ | Dissociation rate of Pol-II in the absence of crowders | 1nM |
| $\boldsymbol{k}_{\boldsymbol{m}}$ | Transcription rate of Pol-II in the absence of crowders | $0.001$s^-1^ |
| $\boldsymbol{r}_{\boldsymbol{min}}$ | Lower length scale of chromatin self-similarity | 1nm |
| $\boldsymbol{\sigma}^{\mathbf{2}}$ | Variance of continues crowding density $\phi$ | $\phi_{in,0}(1-\phi_{in,0})$ |
| **L** | Average number of base pairs in each gene | 6Kbp |
| $\boldsymbol{r}_{\boldsymbol{in}}^{\mathbf{0}}$ | Radius of interaction volume for single base pair | 15nm |
| $\boldsymbol{M}_{\boldsymbol{f}}$ | Total mass of upper length scale of chromatin self-similarity | 10Kbp |
| $\boldsymbol{\phi}_{\boldsymbol{in}\mathbf{,0}}$ | Average crowding density | Chromatin contribution  (BJ cells, ChromEM CVC measurements): 32% v/v  Mobile crowding  contribution: 10% v/v  Total: 42% v/v |
| **Unfixed Parameters** | **DESCRPTION** | **VALUE** |
| ${\mathbf{[}\boldsymbol{C}\mathbf{]}}_{\boldsymbol{tot}}$ | Total concentration of transcription complexes | [0.01nM, 0.1nM] |

**Table S1. Descriptions and values of CPMC model parameters**

**SI Method**

**Calculation of Chromatin Packing Scaling (*D*) from spectral variance (**$\Sigma^{2}$**) measured by PWS**

***Mathematical description of chromatin within the fractal regime by autocorrelation function***

As suggested by the ChromSTEM experiment, chromatin is likely to form a fractal structure within packing domains (PDs). Within the fractal regime, the genomic size of the chromatin ($N_{f}$) scales up with its physical size$r_{f}$ following a power-law relationship (*1*).

$N_{f}= \left( \frac{r_{f}}{l_{min}} \right)^{D}$	1

where *D* is the chromatin packing density scaling (fractal dimension), $l_{min}$ ~ 1 nm is the radius of the fundamental structural unit of chromatin, the nucleotide base pair, $r_{f}$ is the upper bound of the power-law regime which represents the radius of the PD.

The autocorrelation function (ACF) representing chromatin mass density within the fractal regime adopts a power-law function, with the same power exponent *D*. However, a strict power-law function approaches infinity at the origin, a behavior that is not physical, as the smallest structural unit of chromatin are nucleotides which have finite size. Additionally, the true ACF of a single PD gradually decays to zero. Thus, a modified power-law ACF ($B_{\rho}$) was used to include a lower and upper length limit to the power-law regime, and allow for both continuity and differentiability for all length scales (*2*, *3*), as shown in equation [2]:

$B_{\rho}\left( r,D_{B}, l_{min},l_{max} \right)= \sigma_{\rho}^{2} \frac{D_{B}-3}{{l_{max}}^{D_{B}-3} -{l_{min}}^{D_{B}-3}} r^{D_{B}-3}\left[ \Gamma\left( \frac{r}{l_{max}}, 3-D_{B} \right)- \Gamma\left( \frac{r}{l_{min}}, 3-D_{B} \right) \right]$	2

where $r$ is the spatial separation, $\Gamma(x, a)$ is the upper incomplete gamma function, and $l_{min}$ and $l_{max}$ characterize the lower and upper length scales of fractality, respectively. $D_{B}$ is the effective chromatin packing scaling, a model parameter that describes the shape of $B_{\rho}$ and is related to *D*, the true chromatin packing scaling. $\sigma_{\rho}^{2}$ $\frac{D_{B}-3}{{l_{max}}^{D_{B}-3} -{l_{min}}^{D_{B}-3}}$ is the normalization factor of this ACF model, such that $B_{\rho}$(r = 0) is $\sigma_{\rho}^{2}$, the variance of mass density. We tested the validity of $B_{\rho}$ to represent the ACF of chromatin mass density by fitting the model to experimentally derived ACFs from ChromTEM (50nm sections). For both A549 and BJ cells, the modified ACFs match the experimental ACFs with marginal errors (median R^2^ of 0.985 over all samples for the fitting range of r between 50-200nm) (**Fig. S12)**, demonstrating the flexibility of this model.

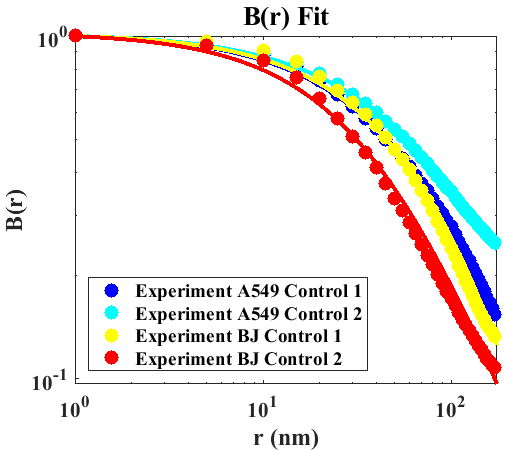

**Fig. S12.** Comparing $B_{\rho}$(r) to representative experimental ACF measured by ChromTEM for A549 cells and BJ cells. For these four examples, the normalized model, shown as solid lines, fits the experimental data with only a small margin of error.

Because of the chosen model, $B_{\rho}$ it is not purely power law up to $l_{max}$ but rather to some distance smaller than $l_{max}$. To account for this, we calculate the input model parameter $l_{max}$ using the following formula to correct for the desired power law maximum point, $l_{\max\_pl}$:

$l_{max}= \left[ \frac{N_{f}D_{B}\left[ 1-\left( \frac{l_{min}}{l_{\max\_pl}} \right)^{{3-D}_{B}} \right]}{\left( 3-D_{B} \right)\left[ 1-\left( \frac{l_{min}}{l_{\max\_pl}} \right)^{D_{B}} \right]} \right]^{\frac{1}{D_{B}}}$	3

In order to numerically establish the relationship between $D_{B}$ and *D*, for each $D_{B}$, we generated $B_{\rho}$ by computing $l_{max}$ from [3]: choosing physiological values of $N_{f}$, setting $l_{min}=1nm$, and starting with an estimate of $l_{\max\_pl}= l_{min}{N_{f}}^{\frac{1}{D_{B}}}$. Then, using linear regression, we fit $B_{\rho}$ to $r^{D-3}$within the range of $\frac{l_{max}+l_{min}}{100}$to $\frac{l_{max}+l_{min}}{100}+0.1$ to compute *D* (**Fig. S13**). Importantly, while $D_{B}$ can range from 1 to 4, *D* can only physically take on values between 5/3 and 3 for a topologically unconstrained polymer in thermodynamic equilibrium. We notice that $D_{B}\approx D$ for all physiological values of *D* except for *D* approaching 3.

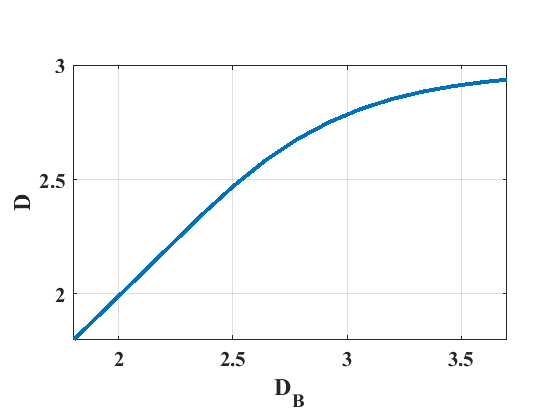

**Fig. S13.** The numerically computed relationship between *D* and $D_{B}$ for the $B_{\rho}$ form described in equation [1].

***Calculating chromatin packing scaling D from PWS signal*** $\Sigma^{2}$

Within the nucleus, chromatin is the major contributor to PWS signal as most other macromolecules and physicochemical elements (i.e. ions) that also comprise the nuclear environment are below the length-scale sensitivity of PWS, which is 20-300nm, as the size of proteins are usually on the order of a few nanometers(*4*). In order to establish a direct relationship between the chromatin packing scaling *D* from PWS spectral variance $\Sigma^{2}$, we first express $\Sigma^{2}$as a function of the effective chromatin packing scaling $D_{B}$, then numerically convert to *D*. Next, we utilize the relationship between spectral variance and the spatial ACF: $\Sigma^{2}\propto\left[ B_{\rho}\left( r \right)\otimes T\left( r \right) \right]|_{r=0}$, denoted by the convolution ($\otimes$) between the ACF and a smoothing function (*T*) characterized by the microscope’s NA and the source spectrum(*5*), evaluated at r = 0. It is clear that $\Sigma^{2}$ is linearly related to the ACF, and a linear decomposition of the ACF would result in a linear addition of $\Sigma^{2}$.

Employing the Laplace transform, we expanded the modified power-law ACF$B_{\rho}\left( r,D_{B}, l_{min},l_{max} \right)$ to a sum of weighted exponential functions, each with a characteristic decay length within the fractal regime [4]

$B_{\rho}\left( r \right)=\int_{l_{min}}^{l_{max}} P\left( l_{c},D_{B} \right)e^{-r/l_{c}} dl_{c}$	4

Here, $e^{-r/l_{c}}$ is a series of exponential basis functions with varying $l_{c}$, the characteristic length that modulates the speed of decay. *P* contains the weights for each exponential basis in the form of a continuous probability distribution function. *P* can be obtained by the normalized inverse Laplace transform. Putting equation [4] into the following form: $B_{\rho}(r)=\int_{0}^{\infty} f\left( t \right) e^{-rt}dt$, allows the calculation of *P* using the inverse Laplace through a simple change of variable, as shown in equation [5]. Furthermore, *P* must be normalized to sum to unity within the fractal regime:

$P\left( l_{c},D_{B} \right)=\frac{{f\left( \frac{1}{t} \right)}/{t^{2}}}{\int_{1/{l_{min}}}^{1/{l_{max}}} {f\left( \frac{1}{t} \right)}/{t^{2}}dt} = {l_{c}}^{D_{B}-4} \frac{D_{B}-3}{({l_{max}}^{D_{B}-3} -{l_{min}}^{D_{B}-3})}$	5

Governed by the unitarity of the Fourier transform from spectral to spatial variance (Parseval's theorem), we can rewrite equation [4] in the spectral space while maintaining the same weighting function *P*. From [4], replacing $B_{\rho}\left( r \right)$ with $\Sigma^{2}(D_{B})$ and $e^{-r/l_{c}}$ with $\Sigma_{e}^{2}(l_{c})$, the spectral variance measured from an exponential ACF, we obtain the relationship between PWS signal $\Sigma^{2}$ and the effective chromatin packing scaling $D_{B}$:

$\Sigma^{2}(D_{B})=\int_{l_{min}}^{l_{max}} P\left( l_{c},D_{B} \right) \Sigma_{e}^{2}(l_{c}) {dl}_{c}$	6

Considering the experimental setup of a PWS microscope (Materials and Methods: PWS imaging), $\Sigma_{e}^{2}(l_{c})$ has a closed-form solution for an exponential basis function with characteristic length $l_{c}$(*6*):

$\Sigma_{e}^{2}(l_{c})= \frac{2R^{2}\sigma_{n_{\Delta}}^{2}}{\pi}\frac{{l_{c}^{3}k}^{4}L{NA}^{2}}{[1+ k^{2}l_{c}^{2}(4+{NA}^{2})](1+4k^{2}l_{c}^{2})}$	7

In [7], *R* is the product of the forward and reverse Fresnel transmission and reflection coefficients at the cell/glass interface, normalized by the reflectance coefficient of the reference (glass/media) interface: $\frac{4 n_{nucleus} n_{glass}(n_{glass} - n_{nucleus})}{{({n_{glass} + n}_{nucleus})}^{3}}\frac{{(n_{glass} + n_{media})}^{2}}{{(n_{glass} - n_{media})}^{2}}$, *k* is the center wavelength in vacuum, *NA* is the collection numerical aperture in air, *L* is the effective thickness of the sample, determined by the minimum of either the optical cell thickness or the depth of field (DOF), and $\sigma_{n}^{2}$ is the variance of refractive index (RI) fluctuations within the nucleus .

Next, we estimated the RI of the nucleus $n_{nucleus}$ from the densities of these nuclear components through the Gladstone-Dale equation:

$n_{nucleus}\left( \phi\right)=n_{0}+ \alpha\rho_{chromatin}\phi+ \alpha\rho_{MC}\phi_{MC}$	8

where $n_{0}$ is the RI of water in the wavelength range used; $\alpha= 0.18 \frac{{cm}^{3}}{g}$ is the RI increment and is constant for all macromolecules that contribute to the spectral signal (*7*); $\rho_{chromatin}$ and $\rho_{MC}$ are the densities of chromatin and MCs, $\phi$ is the crowding density of chromatin, and $\phi_{MC}$ is the crowding density of MCs. As most of the MCs we consider are proteins and nucleic acids, we used $\rho_{MC}$ = 1.25 $\frac{g}{{cm}^{3}}$, the average density of pure, dehydrated proteins(*8*). We further estimated $\rho_{chromatin}$to be 0.555 $\frac{g}{{cm}^{3}}$ by approximating the weight and total volume occupied by a single nucleosome and its linker DNA. We inputted a series of $\phi$ between 0.12 and 0.55, within the physiological range of chromatin volume concentration reported by ChromEMT for interphase nuclei (*9*). Finally, we estimate $\phi_{MC} = \phi_{{MC}_{max}}(1-\phi)$, where $\phi_{{MC}_{max}}$ = 0.05 and is the maximum concentration occupied by MCs, and thus $\phi_{MC}$ is proportional to the volume unoccupied by chromatin.

We estimated the standard deviation of RI fluctuations $\sigma_{n}$ by assuming $\phi$ follows a binomial distribution:

$\sigma_{n}= \sqrt{\phi\left( 1- \phi\right)} \left[ n_{nucleus}\left( \phi=1 \right) - n_{nucleus}\left( \phi=0 \right) \right]$	9

We numerically calculated a series of $\Sigma(D_{B})$ for varying $D_{B}$ by inputting physiologically relevant values for $\phi$ and $N_{f}$, and computing $n_{nucleus}$, and $\sigma_{n}$ from equations [8], [9] , respectively. Next, we input these values into equations [5] and [7] to compute $P$ and $\Sigma_{e}^{2}$ as inputs to equation [6]. Importantly, the relationship between $\Sigma$ and $D_{B}$ can be accurately represented by a linear approximation. For physiologically relevant values ($\phi=0.32$, $N_{f}=1.0$Mbp), we obtain

$\Sigma\left( D_{B} \right) \approx A(D_{B}-D_{0})$,	10

with fitted values of A = 0.13, $D_{0}=$ 1.46, and corresponding R^2^ = 0.999, although this model does not change significantly for other physiologically relevant values of $N_{f}$ and $\phi$.

Finally, we scanned through an exhaustive range of possible *D* values and inputted the system incident *NA* of 0.55 and collection *NA* of 1.49 to generate a numerical relationship describing $\Sigma(D)$ as a function of $\phi$ and $N_{f}$. The range of $N_{f}$ values displayed encapsulates the extreme values for packing domain size measured by ChromSTEM, which we have shown exhibits fractal behavior. For calculations performed in calculating D from $\Sigma$ in the main text, we used $\phi$ = 0.32 and $N_{f}$ = 1.0Mbp. As evident in Fig. S14, the inversion, allowing for calculation of *D* given experimentally measured $\Sigma$, is possible due to the monotonicity of the relationship. We also note the primary contributor to changes in $\Sigma$ are changes in *D*, even considering the extreme limits of physiologically relevant $\phi$ and $N_{f}$*.*

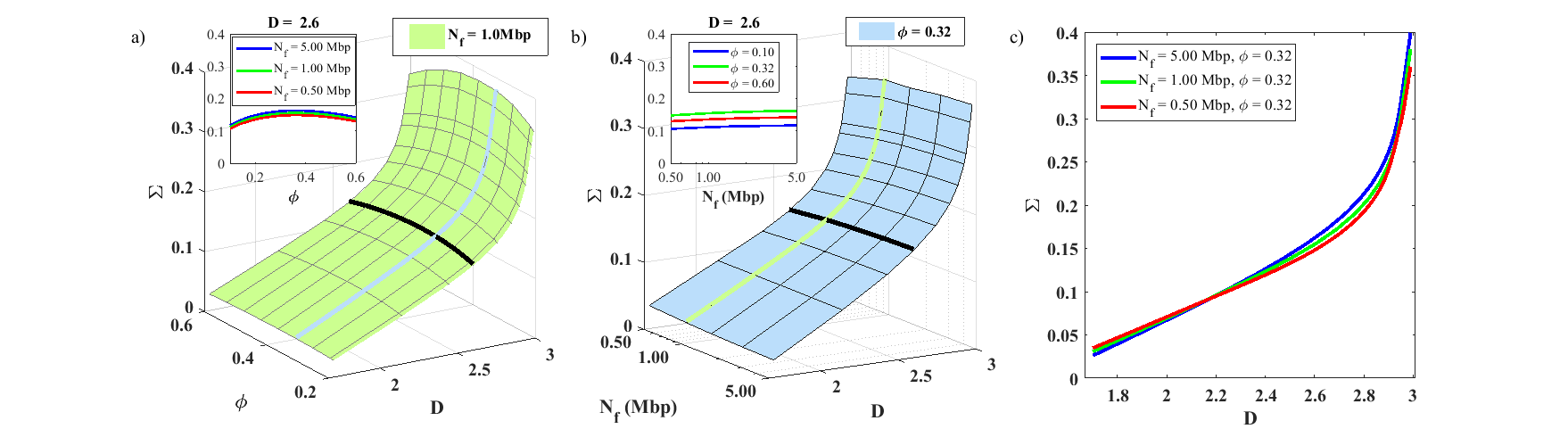

**Fig. S14.** $\Sigma$ **vs. *D,***$N_{f}$ and$\phi$ a-c. Surface plot showing $\Sigma$ vs *D* and $\phi$ for a fixed $N_{f}$ of 1.0Mbp and under varying physiologically relevant conditions and fixed D = 2.6 for $N_{f}$ (denoted by black line and expanded in inset). Surface plot showing $\Sigma$ vs *D* and $N_{f}$ for $\phi$ of 0.32, and for varying physiologically relevant values of $\phi$ for a fixed D = 2.6 (denoted by black line and expanded in inset). In (c), blue and red lines are plotted at the extreme ranges of physiologically relevant values for $N_{f}$, while green shows mapping used in calculations for this article.

1.	T. G. Dewey, *Fractals in Molecular Biophysics.* (Oxford University Press, 1997).

2.	M. Xu, R. R. Alfano, Fractal mechanisms of light scattering in biological tissue and cells. *Opt. Lett.* **30**, 3051–3 (2005).

3.	C. J. R. Sheppard, Fractal model of light scattering in biological tissue and cells. *Opt. Lett.* **32**, 142 (2007).

4.	R. Phillips, J. Kondev, J. Theriot, *Physical Biology of the Cell* (Garland Science, 2009).
